# Pipeline Olympics: continuable benchmarking of computational workflows for DNA methylation sequencing data against an experimental gold standard

**DOI:** 10.1101/2024.09.16.609142

**Authors:** Yu-Yu Lin, Kersten Breuer, Dieter Weichenhan, Pascal Lafrenz, Agata Wilk, Marina Chepeleva, Oliver Mücke, Maximilian Schönung, Franziska Petermann, Philipp Kensche, Lena Weiser, Frank Thommen, Gideon Giacomelli, Karl Nordstroem, Edahi Gonzalez-Avalos, Angelika Merkel, Helene Kretzmer, Jonas Fischer, Stephen Krämer, Murat Iskar, Stephan Wolf, Ivo Buchhalter, Manel Esteller, Chris Lawerenz, Sven Twardziok, Marc Zapatka, Volker Hovestadt, Matthias Schlesner, Marcel Schulz, Steve Hoffmann, Clarissa Gerhauser, Jörn Walter, Mark Hartmann, Daniel B. Lipka, Yassen Assenov, Christoph Bock, Christoph Plass, Reka Toth, Pavlo Lutsik

## Abstract

DNA methylation is a widely studied epigenetic mark and a powerful biomarker of cell type, age, environmental exposures, and disease. Whole-genome sequencing following selective conversion of unmethylated cytosines into thymines via bisulfite treatment or enzymatic methods remains the reference method for DNA methylation profiling genome-wide. While numerous software tools facilitate processing of DNA methylation sequencing reads, a comprehensive benchmarking study has been lacking thus far. In this study, we systematically compared complete computational workflows for processing DNA methylation sequencing data using a dedicated benchmarking dataset generated with five genome-wide profiling protocols. As an evaluation reference, we employed highly quantitative locus-specific measurements from our preceding benchmark of targeted DNA methylation assays. Based on this experimental gold-standard assessment and several comprehensive metrics, we identified workflows that consistently demonstrated superior performance and revealed major workflow development trends. To facilitate the sustainability of our benchmark, we implemented an interactive workflow execution and data presentation platform, adaptable to user-defined criteria and seamlessly expandable to future software.

## Introduction

DNA methylation is a key epigenetic modification [1] that plays essential role in development [2] and cell differentiation [3, 4] across many species including human. DNA methylation landscapes are altered in the course of mitotic divisions [5] and transition to cellular senescence [6], during ageing [7, 8], as well as in pathological conditions including cancer [9–13] and other diseases [14, 15]. The high stability of DNA methylation compared to gene expression and the ease of analysis compared to other epigenomic marks contribute to the attractiveness of DNA methylation as an epigenetic biomarker of age [16], for early detection of cancer in liquid biopsies [17], and in forensic assays [18].

In eukaryotes, DNA methylation occurs predominantly at CpG dinucleotides. Numerous methods have been proposed to measure CpG methylation patterns, as reviewed [19, 20] and evaluated in primary research papers [21–25]. The most comprehensive is whole-genome bisulfite sequencing (WGBS), which provides a genome-wide, single-base pair and single-strand resolution method based on the bisulfite conversion of unmethylated cytosines [26]. The Illumina Infinium microarrays [27, 28] as well as reduced representation bisulfite sequencing [29] provide additional genome-scale alternatives, measuring 2-15% of the CpG sites. These methods have to be distinguished from focused assays, such as amplicon bisulfite sequencing and bisulfite pyrosequencing [21].

Bisulfite treatment results in the chemical deamination of unmethylated cytosines and subsequently their change to thymines. This induces DNA fragmentation and degradation, thus requiring high amounts of DNA input [24]. To overcome this issue, a variety of enhanced protocol variants for moderate to low input DNA amounts have been suggested, including tagmentation-based WGBS (T-WGBS) [30–32] and post-bisulfite adaptor tagging (PBAT) [33]. The former increases the efficiency of standard adaptor tagging, whereas the latter utilizes post-bisulfite adaptor tagging to avoid subsequent degradation of adaptor-tagged fragments. Finally, enzymatic methods such as EM-seq replace bisulfite treatment with an enzymatic conversion step, which reduces DNA fragmentation and degradation [34]. Most bisulfite sequencing protocols currently in use are directional, whereby the actual sequencing reads correspond to a converted version of either the original forward or reverse strand (OT and OB strands). Depending on the design of adaptor ligation, non-directional protocols exist, such as PBAT, whereby all four possible converted DNA strands are sequenced at roughly the same frequency [32]. All these protocol-specific differences require special attention during data processing.

Analysis of bisulfite sequencing data generally includes four core steps: 1) read processing, including quality control and trimming; 2) conversion-aware alignment; 3) post-alignment processing or filtering; and 4) calling of methylation states (and, optionally, of genotypes and structural variation). Many tools have been developed for each step, providing room for an overwhelming number of possible combinations and workflows. Read preprocessing includes basic QC [35], standard read trimmers [36, 37], bisulfite alignment, alignment post-processing, as well as methylation calling and quantification. Methods to account for bisulfite conversion during the alignment include no-cytosine three-letter alphabet [38–44], a wild-card alignment [45–48], or a wild-card-related approach that transforms the alignment into an asymmetric mapping problem [49]. In the three-letter approach, all cytosines in both the reference genome and the sequencing reads are converted to thymines, and mapped using a seed and extend approach [50]. Wildcard aligners map both cytosines and thymines in the reads to cytosines in the reference genome. Post-processing includes filtering PCR duplications with conventional tools as well as other quality filtering steps, for example, filtering by alignment quality. Finally, calling and quantification of methylation states range from simple counting [51, 52] to Bayesian model-based approaches [53] featuring local realignment [54]. Some methylation callers provide add-on functionalities, such as sequence variant calling [53–56].

Although numerous methods of profiling methylation have been proposed, no comprehensive attempt has been made at evaluating end-to-end data processing workflows. Previous benchmarks focused on a single processing task (typically the alignment), assessed relatively few methods and lacked gold-standard control datasets [57–59].

Here we present a comprehensive benchmark of data processing workflows for DNA methylation sequencing. Our study is based on gold-standard samples with highly accurate DNA methylation calls. We evaluated the workflows in the context of two standard and three low-input Methyl-Seq protocols. To simplify the choice of workflows for the users and enable seamless extension to future tools and workflows, we developed web-resources for interactive data presentation and continuous and long-lasting (“living”) benchmarking.

## Results

### Systematic review and selection of benchmarked software and workflows

We conducted a comprehensive literature search and reviewed published software tools for bisulfite sequencing data processing (Supplementary Table 1). We focused on complete workflows covering processing from raw reads to DNA methylation calls and excluded those that were not open-source or not regularly maintained (see Methods for details). Altogether, we included 10 methylation calling workflows into our study: *BAT* [60], *Biscuit* [61], *Bismark* [40], *BSBolt* [62]*, bwa*-*meth* [51], *FAME* [49], *gemBS* [53], *GSNAP* [63], *methylCtools* [52], and *methylpy* [64] (**Figure 1a**). With this selection, we cover different approaches to bisulfite alignment (3-letter and wild-card) and DNA methylation calling (counting and Bayesian-based).

**Figure 1:**
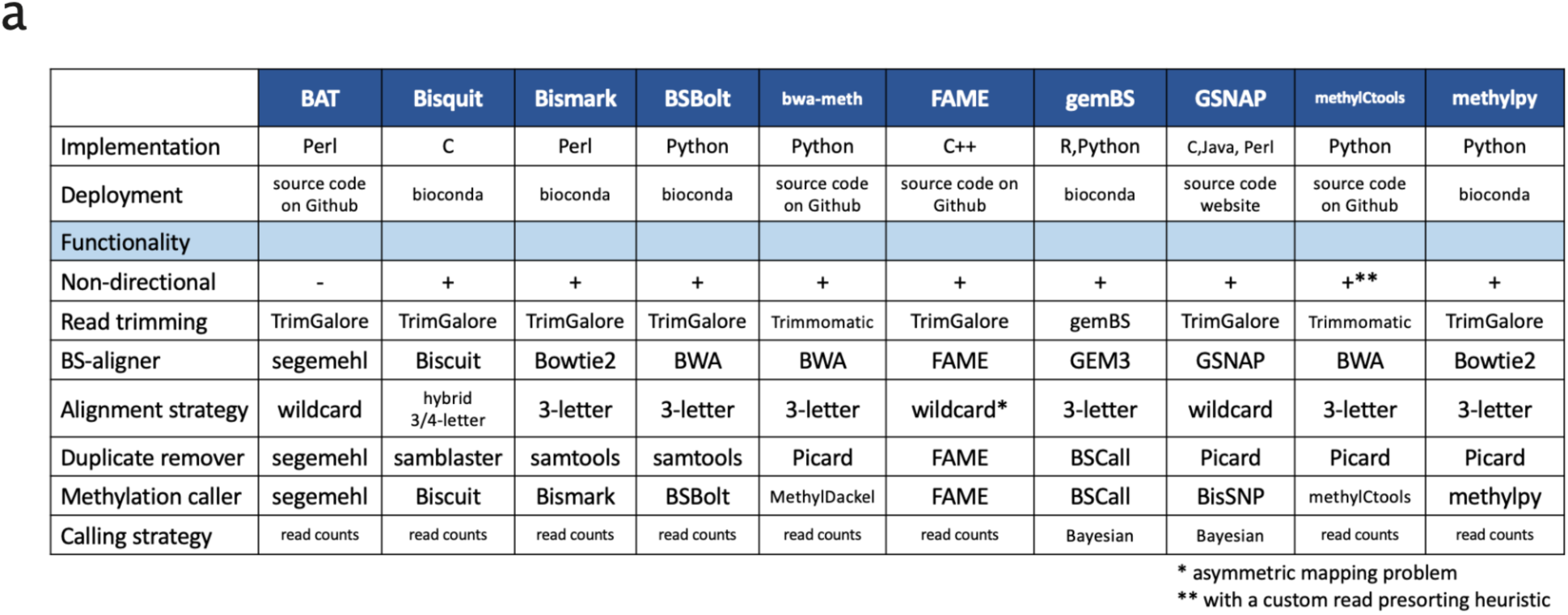
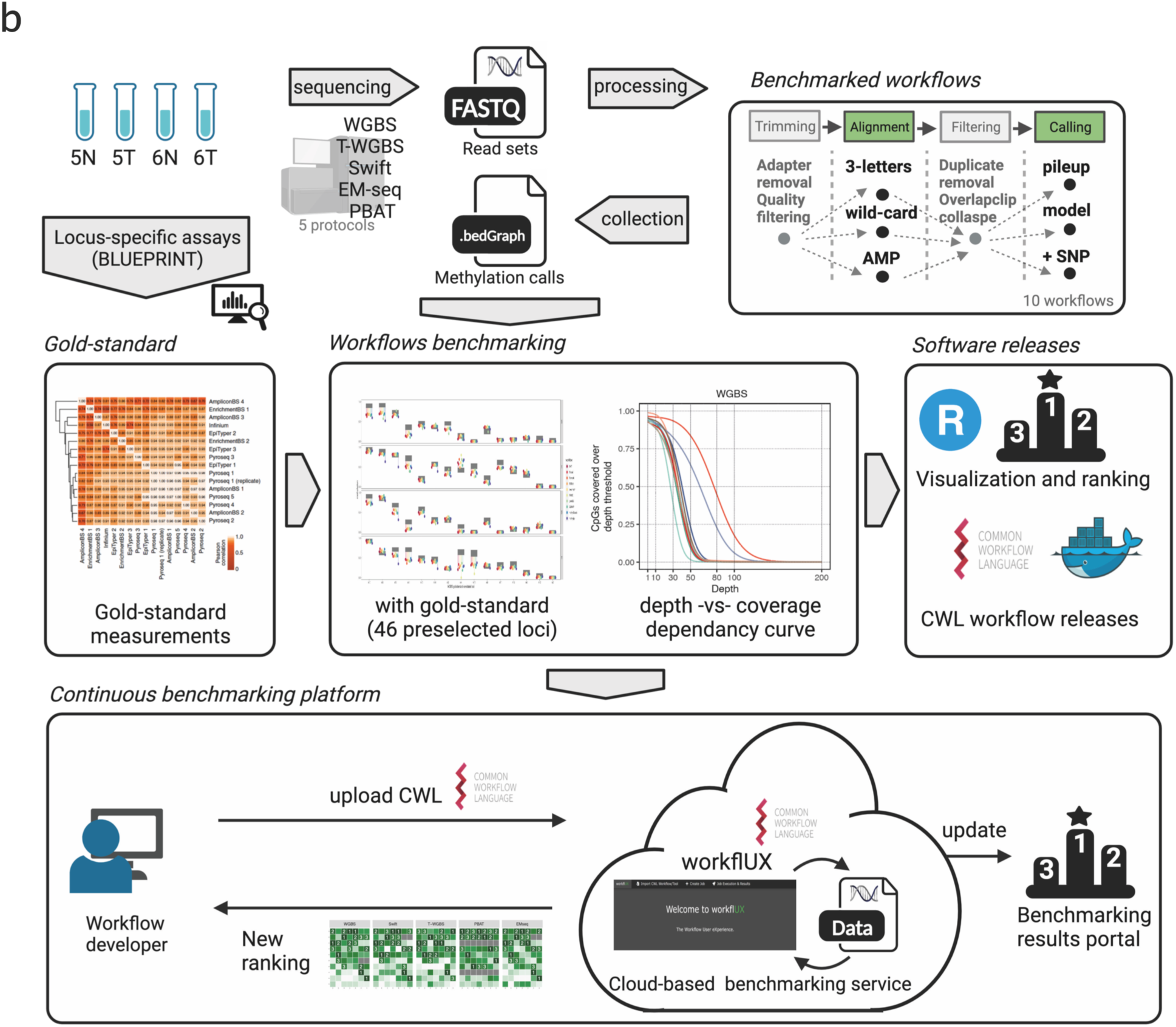
Workflow selection and study design. **a.** The composition and main characteristics of the evaluated workflows. The criteria for choosing the workflows are explained in the Methods section, under section ‘Workflow Selection and Installation”. **b**. Schematic overview of the study design. Two pairs of colon cancer and adjacent normal samples were selected from [21] featuring consensus methylation corridors from multiple high-resolution methods at selected loci that served as the “gold-standard” measurements for our project. All samples were sequenced using five different methyl-sequencing protocols, including one standard (WGBS, SWIFT), two low-input (T-WGBS, PBAT) and one bisulfite-free (EM-seq), and the data were processed using ten selected workflows. Common Workflow Language (CWL) workflows and a Shiny-based portal for visualization are provided. To support the development of future workflows, we introduce a dedicated workfIUX server to allow developers to execute their workflow on the selected datasets and compare the result with those presented in this study. The new rankings will be updated on the open benchmarking service.

### Benchmarking study design and dataset

We selected two tumor-normal pairs of colon cancer originally used for benchmarking locus-specific methylation profiling technologies by the BLUEPRINT consortium [21] (**Figure 1b**). In this study, a few preselected genomic regions were profiled in a range of samples, including six colon tumor / adjacent normal sample pairs, and multiple labs performed 16 targeted DNA methylation assays, including AmpliconBS, EnrichmentBS, EpiTyper, Infinium, and Pyroseq. The combination of multiple targeted arrays and the collaboration of multiple labs resulted in highly accurate DNA methylation calls. This study established consensus corridors for the true DNA methylation levels at the assayed sites, which we use here as the gold-standard loci set for benchmarking. We sequenced all four samples using a representative set of five Methyl-Seq protocols: two standard (WGBS, SWIFT), two low-input (T-WGBS, PBAT), and a bisulfite-free enzymatic protocol (EM-seq) (Supplementary Figure 1). We obtained high-quality raw sequencing data ranging from 300,312,952 (PBAT, 6T) to 985,933,822 (WGBS, 6T) read pairs per sample (Supplementary Table 2). We implemented all selected workflows in CWL (Common Workflow Language [65]) and ran them on a dedicated virtual machine in a fully controlled computational environment. The resulting methylation calls were aggregated and summarized using *methrix* [66] and evaluated using a range of criteria to establish a consistent ranking. Finally, we set up a cloud-based infrastructure to establish an environment that will allow researchers the continuation of benchmarking using their own workflows.

To understand the challenges each library preparation protocol poses for processing workflows, we first assessed the major properties of the generated data. Thus, we examined the data emerging from each protocol to assess the extent of variation between sequencing protocols that might affect data quality and processing. We used median measurements from all workflows for visualization. All protocols generated high-quality reads, and quality trimming led to the loss of less than 2% of the data. Alignment rates were at least 92%, except for PBAT, where it dropped to 74%. Between 13% and 28% of the aligned reads were identified as PCR duplicates and removed (Supplementary Figure 3, Supplementary Table 3). As expected, the protocols showed significant variation in genome coverage, context preference, and DNA methylation bias. WGBS exhibited the highest depth of CpG coverage with a median of > 43 reads for all samples, whereas the low-input protocols, T-WGBS and PBAT, reached a median coverage of less than 13 reads (**Figure 2a**, Supplementary Figure 2a and Supplementary Table 3). The sequencing depth was relatively uniform across the genome for most protocols and showed a characteristic bell-shaped distribution. In contrast, PBAT showed a positively skewed distribution (**Figure 2a**, Supplementary Figure 2a) and a preference for GC-rich regions (**Figure 2b**, Supplementary Figure 2b). We excluded loci with a sequencing depth below 10 reads and explored the distribution of beta-values, revealing strong variation among different protocols. Each protocol exhibited a broad mode at a beta value of approximately 0.8 (**Figure 2c**). Additionally, we found that, unlike other protocols, WGBS did not exhibit another sharp mode at beta-value=1 corresponding to fully methylated sites. Absence of this peak was due to a systematically higher depth of coverage, as it accentuated when downsampling WGBS data to the genome-wide coverage level of PBAT (Supplementary Figure 4). All protocols showed a protocol-specific methylation ratio shift at the end of the reads (M-bias), which required additional trimming or clipping (Supplementary Figure 2c).

**Figure 2:**
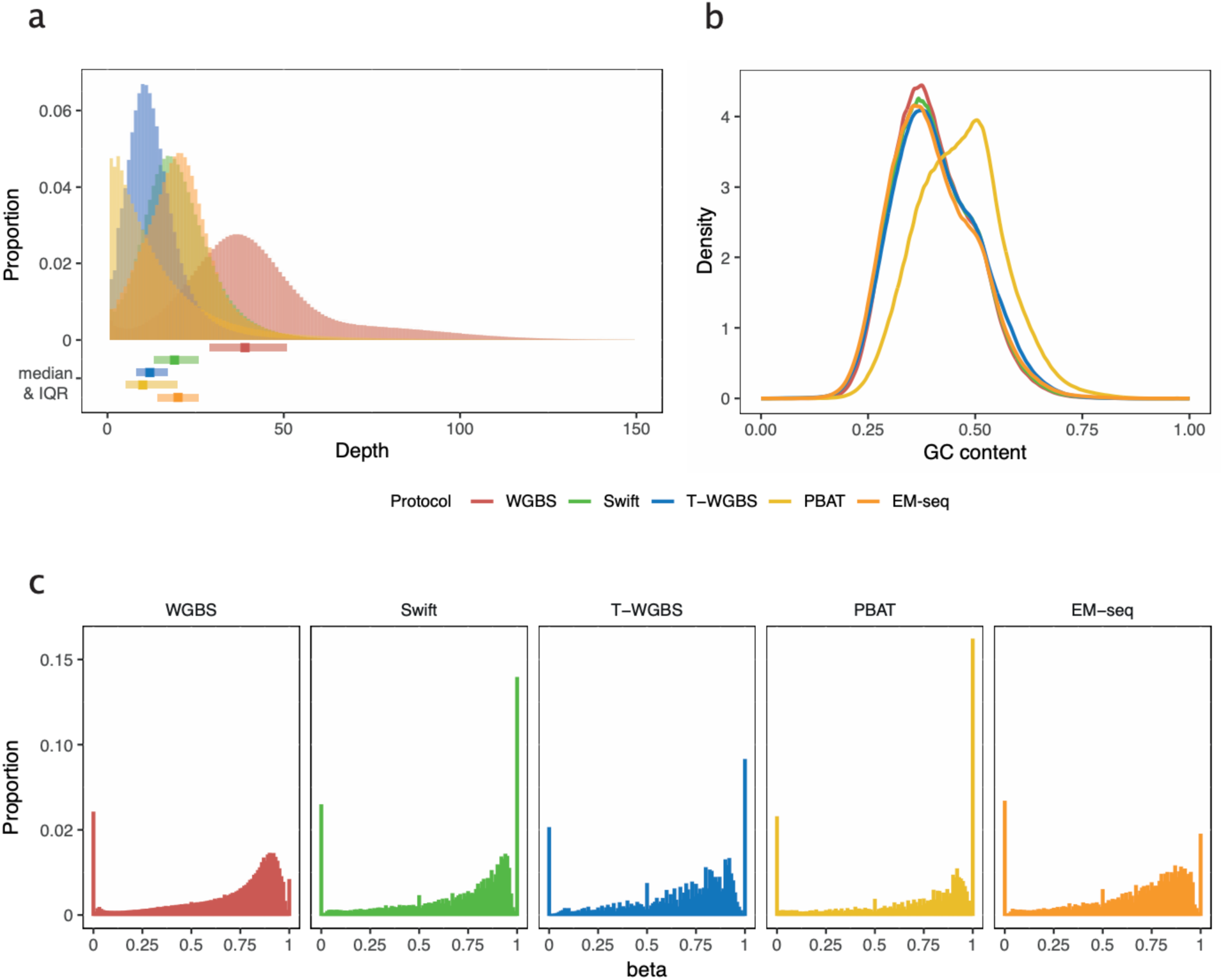
Overview of sequencing protocol differences. **a.** The distribution of coverage depth for the different protocols. The boxplots at the bottom of the histogram indicate the median and interquartile range (IQR) values, offering insight into the central tendency and spread of the coverage distribution. **b.** The distribution density of GC content for aligned reads. Note, that not the read sequence itself but the reference genome sequence at the corresponding position was used for calculation. **c.** The distribution of beta-values resulting from the protocols. CpGs covered by less than 10 reads were excluded. For panels 2a and 2c, we filtered 1% of all CpG sites and used the results of all workflows for the selected CpG sites for visualization.

The single nondirectional protocol in the study, PBAT, poses special challenges for data processing. For instance, it was shown that PBAT library preparation produces so-called chimeric reads containing sequences from two or more distinct genomic loci [48]. Bisulfite treatment and post-PCR adaptor ligation generate four possible sources of reads, and the presence of chimeric reads further complicates this situation, exacerbating the difficulties in alignment. This is primarily attributed to the increased occurrence of multiple best alignment hits, resulting in higher ambiguity and reduced alignment rates. Furthermore, the fragments that were more likely to generate chimeric read pairs were not a random subset of all reads, possibly biasing methylation calling. Therefore, proper handling of chimeric reads not only improves coverage, but also increases accuracy. Thus, we calculated the proportion of chimeric reads relative to the total mapped reads, considering reads that mapped to different chromosomes to be chimeric (Supplementary Figure 5). This allowed us to estimate the extent to which they occurred in different protocols. The results showed that the chimeric read proportion generated in PBAT was 6.64 times that of Swift and 10.33 times that of WGBS.

### Genome-wide analysis of read coverage patterns identifies outlier workflows

We applied the 10 evaluated workflows to the data of the five protocols, resulting in a total of 192 processing runs (*BAT* did not include support the non-directional PBAT protocol and *gemBS* failed on the PBAT data with an unresolvable error). A detailed summary of the data processing steps, with read counts after each step, is given in Supplementary Table 3. The workflows showed similar patterns of alignment rates for all input protocols, except for PBAT. Here, workflows lost 20 - 77% of the reads during alignment, and even the better-performing ones, such as *BSBolt*, had less than 60% of the total reads remaining (Supplementary Figure 3). The fraction of identified PCR duplicates was consistent across the workflows that included this step (Supplementary Table 3).

Although there were significant differences in the protocols and amount of input material used, the majority of the genomic CpG sites were covered with at least one read by most protocol-workflow combinations (**Figure 3a**). We observed greater differences in low-input protocols, such as a lower percentage of CpGs covered in the T-WGBS using the *Biscuit* workflow. PBAT showed the largest discrepancies in the number of covered CpGs between workflows; however, the better performing ones, *BSBolt, Biscuit, FAME* and *methylCtools*, reached genome-wide coverage like other protocols.

**Figure 3:**
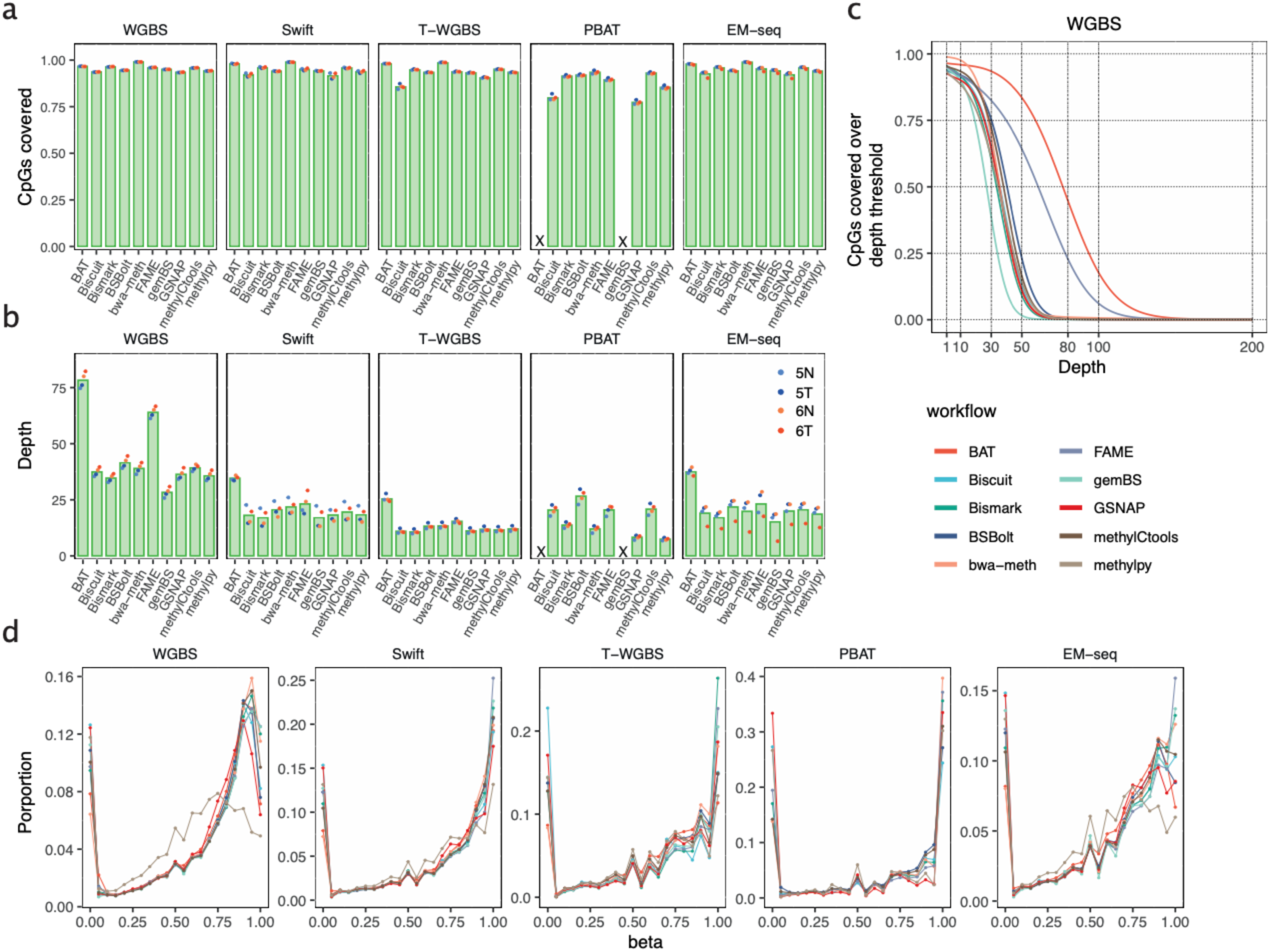
Genome-wide performance and data quality. **a.** The median fraction of total genomic CpGs covered in the output of each workflow by at least one read. Individual samples represented by dots. A cross mark is used to indicate the absence of measurements for the corresponding workflow. **b.** Average number of reads covering the CpG sites, with dots representing the individual samples. A cross mark denotes the absence of measurements in the respective workflow. **c.** The percentage of covered CpGs (y-axis) with read depth below a threshold (x-axis). It provides a practical method for determining the proper depth cut-off threshold. **d.** Distribution of beta-values returned by each evaluated workflow across all experimental protocols. The data points were sampled from all CpGs using a selection rate of 1/100.

We next assessed the depth of read coverage at individual CpGs which is key for accurate estimation of methylation levels (**Figure 3b**). The data revealed that processing with *BAT* and *FAME* generated the highest sequencing depth compared to all other protocols. Substantial variation was detected in PBAT, where the observed variation strongly correlated with the alignment rate (Supplementary Table 3). To ensure the statistical reliability of CpG-wise methylation levels and to make sure that methylation calls have reasonable support for downstream analyses, each CpG site must be covered by multiple reads. A threshold for sequencing depth, selected in practice, is usually traded off against the genome-wide coverage to ensure a large proportion of CpG sites is retained. The depth-vs-coverage dependency curves for each workflow and sequencing protocol (**Figure 3c** and Supplementary Figure 6) revealed that most workflows showed comparable trends, with 75% of CpGs covered with at least 10 reads in WGBS, SWIFT, and EM-seq data (except for *gemBS* in EM-seq). In WGBS, two workflows (*BAT* and *FAME*) showed increased read retention compared to all other workflows (**Figure 3c**). In the case of *BAT*, this can be explained by the lack of duplicate removal. The high coverage in *FAME* data could be attributed to the double counting of CpG calls from the overlapping read mates. It was amplified in WGBS due to the shorter fragment size (**Figure 3c**, Supplementary Figure 4 and Supplementary Table 3). In T-WGBS, the samples had a lower depth, and all workflows, except for *Biscuit*, retained more than 75% of CpGs at the read coverage cut-off of 5. The largest differences between workflows were observed in PBAT, whereby only two workflows, *methylCtools* and *BSbolt,* retained at least 75% of CpGs with a coverage cut-off of 5, whereas two workflows, *methylpy* and *GSNAP*, lost half of the methylation sites with the same cutoff (Supplementary Figure 6). To integrate coverage performance in the final evaluation, we introduced an area-under-curve metric integrating the depth and genome-wide breadth of coverage (see Methods) as a quantitative measure of coverage retention.

Taken together, our evaluation of the global coverage metrics revealed a surprisingly high variation in coverage depth across the workflows, even in the most deeply sequenced WGBS protocol, with several workflows being stark outliers. This highlighted significant differences in read processing, especially related to alignment and PCR duplicate removal, potentially affecting downstream analyses.

### Data-driven genome-wide methylation call consensus corridors elucidate workflow consistency

After evaluating the genome-wide patterns of coverage and CpG retention, we asked how consistent the resulting DNA methylation calls and methylation level estimates were across workflows. We compared genome-wide DNA methylation levels and observed lower (or in one case equal) methylation levels in the tumor samples for all workflows and protocols (Supplementary Table 3), implying that, despite significant differences in effective coverage, all workflows could capture global methylation differences. Generally, the beta value distributions were similar for all workflows except *methylpy*, with larger variations between protocols (**Figure 3d**, Supplementary Figure 7).

To quantitatively assess the similarity of the methylation calls between workflows, we defined a discrepancy score as the mean of absolute pairwise methylation difference between two methylation call vectors (**Figure 4a**). As expected, the workflows showed the highest similarity in high-coverage WGBS protocol, with *methylpy* being a single outlier, and an increase in discrepancy on data from low-input protocols and more shallow sequencing depth, with PBAT showing the largest differences (**Figure 4a**). To compare workflows on a genome-wide level, we introduced a data-driven consensus corridor on the CpG level by taking the smallest range covered by at least five workflows per protocol. We included WGBS, SWIFT, and EM-seq results in the calculations and excluded the two low-input protocols (**Figure 4b**). We ranked the workflows based on the proportion of measurements that fell inside the consensus corridor (**Figure 4c**). We used the sum of genome-wide deviations from the consensus corridor as the primary metric to evaluate the efficacy of genome-wide methylome determination (**Figure 4d**). Protocols with higher coverage exhibited superior performance in terms of high accuracy (resulting in measurements closer to zero) and low variability (with minor variations observed within the protocol). We also examined the deviation concerning genomic annotations and concluded that genic regions exhibited lower deviations than intergenic regions (Supplementary Figure 8).

**Figure 4:**
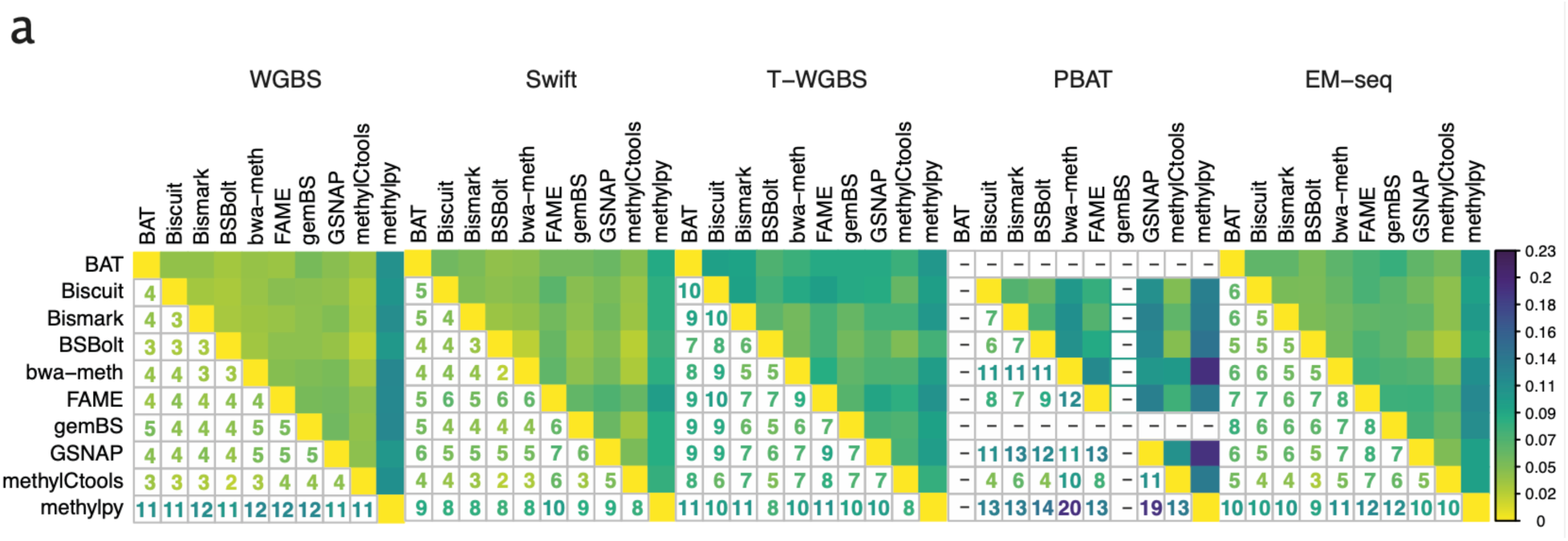
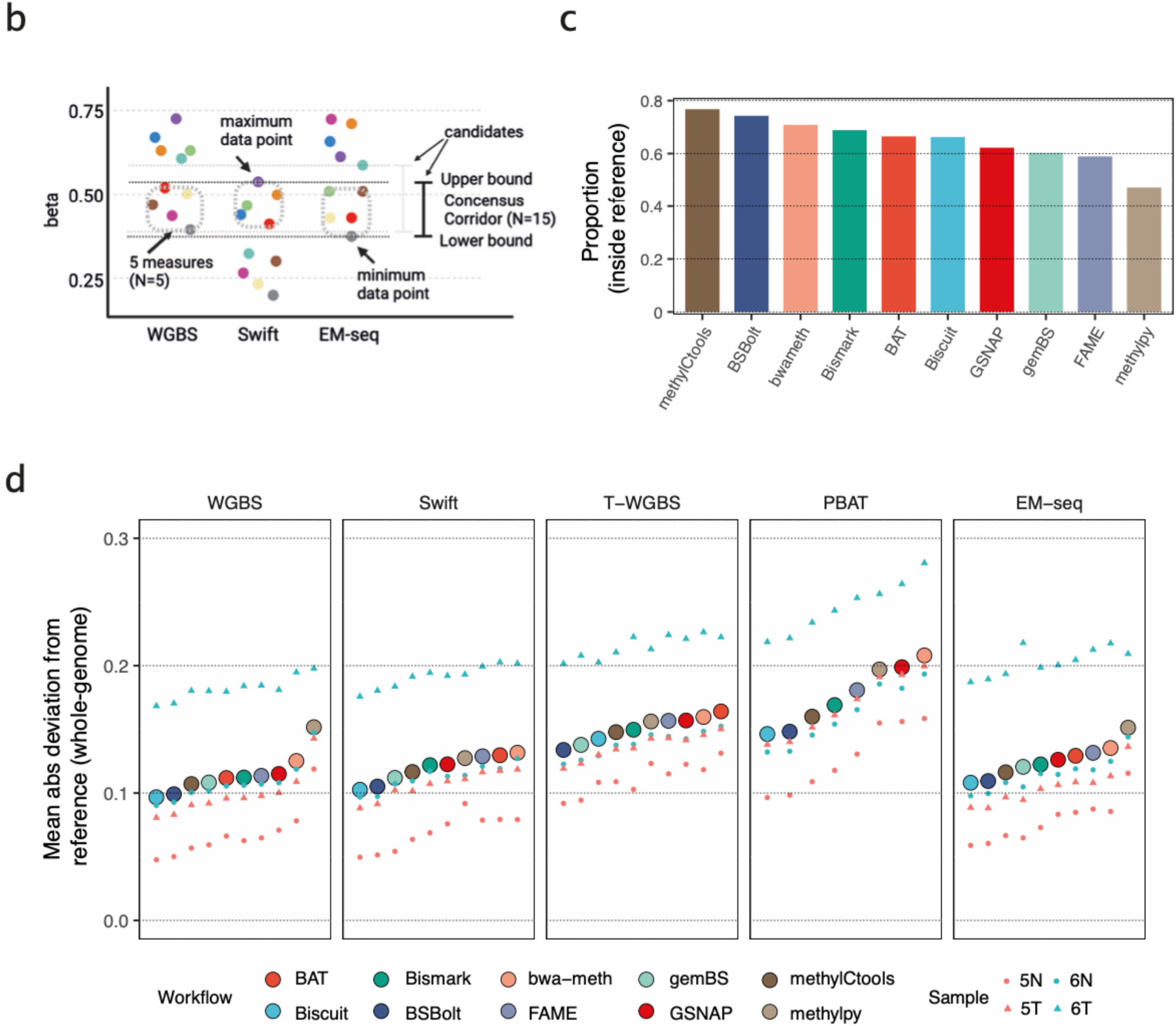
Genome-wide deviation and methylome similarity among workflows. **a.** The methylome dissimilarity between workflows in each protocol. The numbers represent the discrepancy score, defined as mean pairwise difference of beta-values across all CpGs multiplied by 100. Smaller values indicate higher levels of similarity between the methylomes by different workflows. **b.** The definition of genome-wide consensus corridors for all CpGs. The consensus corridor is defined as the smallest region encompassing at least 5 measurements from each of the three high-coverage protocols, WGBS, Swift, and EM-seq. **c**. The fraction of measurements that falls within the all-protocol consensus corridor for each workflow. **d.** Genome-wide mean absolute deviation from the border of the consensus corridors. Within a protocol, the workflows are sorted ascendingly.

In summary, on WGBS, Swift, and EM-seq data the workflows demonstrated better outcomes, with only *methylpy* exhibiting slight outlier trend. Workflow performance on PBAT data showed the highest variability. In this scenario, *Biscuit*, *BSBolt* and *methylCtools* achieved the most favorable results. Notably, all three workflows showed high alignment rates for PBAT data (Supplementary Figure 3) while the ones with higher deviation scores, *bwa-meth* and *methylpy,* were characterized by a much lower alignment rate. This might be explained by workflow differences in handling PBAT chimeric reads. A higher alignment rate may provide a more accurate methylation rate. Indeed, the alignment rate correlated with the deviation score in PBAT (average of sample-wise Spearman = 0.105) (Supplementary Figure 9).

Collectively, the performance of methylation calls between workflows strongly depends on the characteristics of the data and on the specific protocol used to generate it. Technical differences between the workflows did not have major impact upon datasets with higher sequencing depths (WGBS, SWIFT, EM-seq). In contrast, on PBAT data the methylation calls were less consistent, implying that low coverage and technical challenges amplify even minor differences between workflows. Therefore, proper selection of data processing workflows is particularly important for low-input protocols.

### Workflow accuracy evaluation against the experimental reference pinpoints workflow-specific pitfalls

We sought to obtain objective estimates of methylation call accuracy by utilizing the 46 preselected loci from the multi-method and multi-center BLUEPRINT study from which a consensus corridor containing the most likely true methylation values was derived (Supplementary Table 4, see Methods for details) [21]. We used these consensus corridors as ground-truth methylation measurements. Upon initial inspection, we observed that for 45% to 52% of the loci, the methylation values returned by workflows lie within the consensus corridors across all protocols. Certain loci, such as m3 and m4, displayed stronger deviations which were sample-specific and affected all workflows (**Figure 5a**, Supplementary Figure 10 to Supplementary Figure 15).

**Figure 5:**
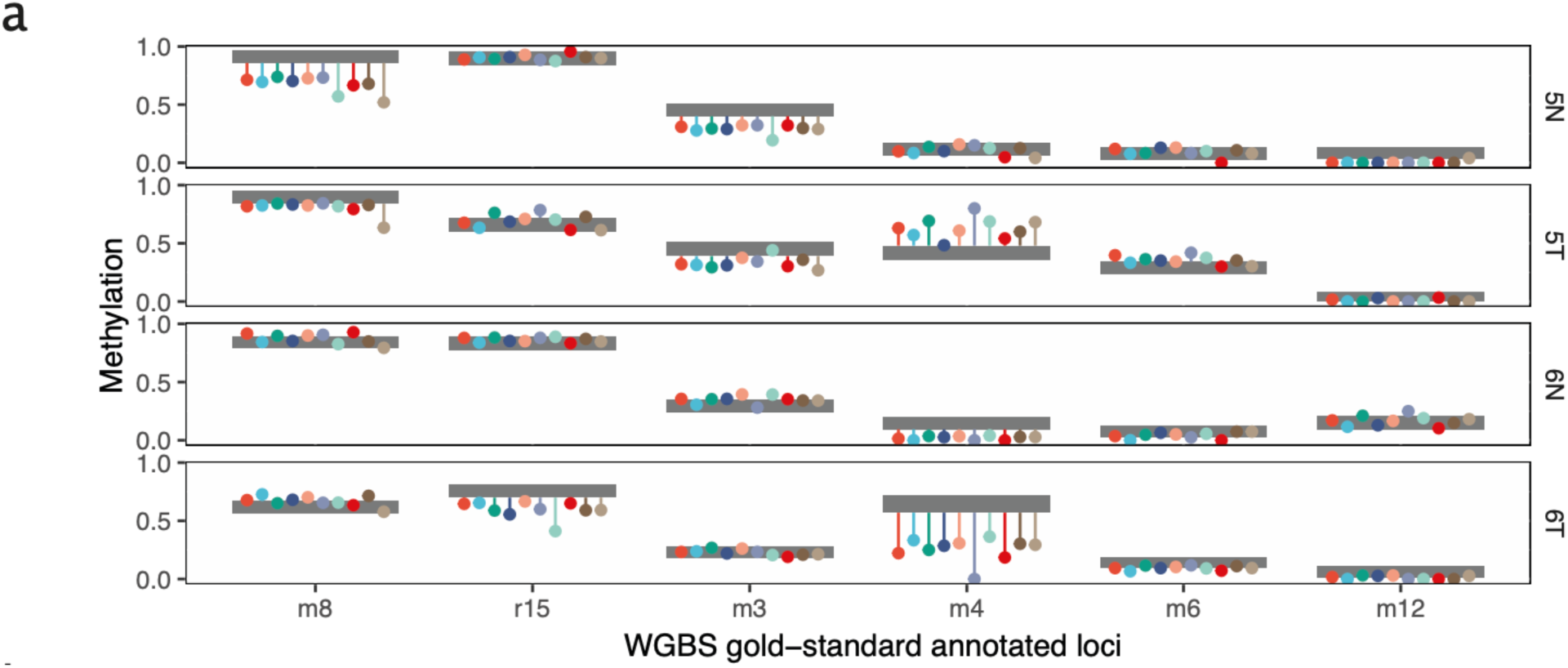
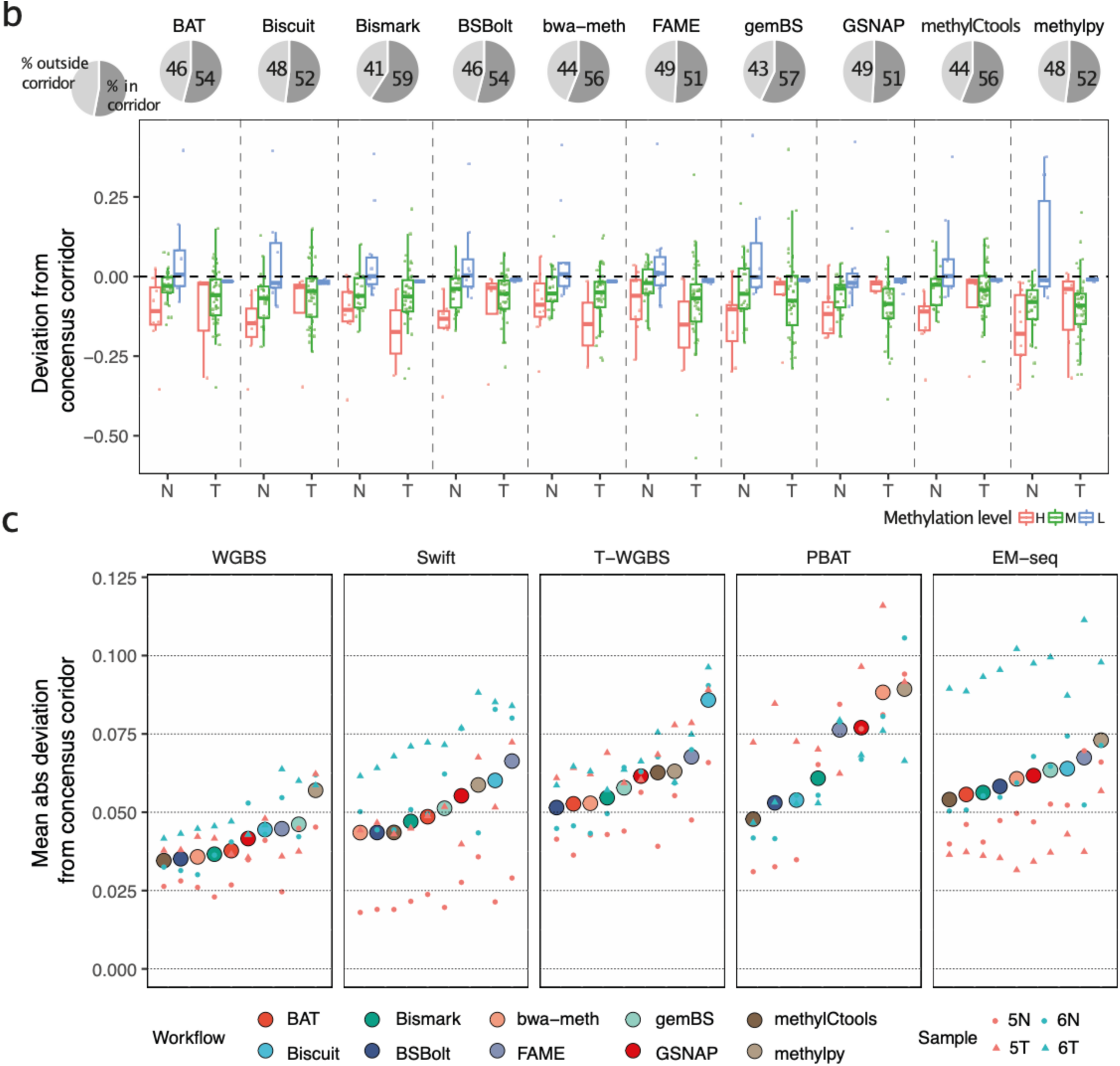
Assessment of methylation call accuracy based on the experimental gold standard. **a.** Deviation from the gold-standard consensus corridors for six selected loci. Several example loci were selected here to represent high (left), intermediate and low (right) methylation levels. Grey boxes represent the consensus corridors (see Methods). The dots show the measured beta values for each workflow, while the lines depict their deviation from the consensus corridors. **b**. Deviation from the consensus corridors of WGBS for all samples combined. The pie charts at the top show the proportion of sites outside/within the consensus corridor. The box plot below shows the distribution of deviation excluding the data inside of the consensus corridor and N/T indicate for normal/tumor respectively. **c.** The mean absolute deviation by protocols and workflows. Within a protocol, the workflows were sorted ascendingly. The deviation is the average of four samples and the deviation of four samples was labeled on the same vertical line.

To summarize the estimates, we calculated the deviation from the consensus corridor at each locus (**Figure 5b**, Supplementary Figure 16). We did not observe strong systematic differences between workflows. Most protocols and workflows, especially WGBS, tended to measure lower methylation levels compared to the consensus measurement across several targeted assays with complementary strengths (**Figure 5b**). On the other hand, the extent of the deviation was highly sample-specific. We observed a tendency that the lowly methylated regions (ß < 0.2) were more accurately measured, whereas the accuracy decreased for the highly methylated (ß >= 0.8) and especially the intermediately methylated (0.2 <= ß < 0.8) regions (**Figure 5b,** Supplementary Figure 16). We calculated the accuracy of the workflow as the mean absolute deviation across the 46 loci (**Figure 5c**). The effect of the workflow was statistically significant only for T-WGBS and PBAT (two-way ANOVA analysis, the p-value for T-WGBS and PBAT are 6.45×10^−4^ and 1.2×10^−3^, respectively, WGBS 0.845, Swift 0.141, EM-seq 0.839) when adjusted for sample-wise differences.

Similarly to the genome-wide consistency analysis, we observed that *methylpy* tended to underestimate methylation at specific loci (**Figure 5b**, Supplementary Figure 16), a pattern inconsistent with other established workflows. We investigated the divergence of *methylpy* from other workflows by using selected gold-standard loci. To illustrate this, aligned reads for *BSBolt* and *methylpy* at locus r23 in the normal sample from Patient 6 were examined using the Integrative Genomics Viewer (IGV) browser. Despite sharing the same alignment, these two tools produced different methylation results (Supplementary Figure 17). Using the Multiple Sequence Alignment (MSA) tool Clustal Omega (https://www.ebi.ac.uk/Tools/msa/clustalo/), we observed that the sequences from the *methylpy* intermediate alignment and the final alignment were different from the raw reads and alignment of *BSBolt* (Supplementary Figure 17). This inconsistency appears to be the result of an error in the simulation of the bisulfite conversion. *Methylpy* employs a three-letter method that conducts *in silico* conversion of reads prior to alignment with the three-letter genomic reference. In the methylation calling step, the converted reads must be restored to their original sequences. Our observations indicated a systematic error or bug at this specific step, resulting in unrestored bases. This provided one explanation for the outlier calls produced by *methylpy*.

Taken together, the analysis of methylation calls at gold-standard loci allowed us to obtain realistic estimates of workflow accuracy and helped us objectively rank the workflows. Furthermore, exemplary deeper investigation of outliers allowed to find specific pitfalls of individual workflows, demonstrating the potential usefulness of our results for workflow debugging.

### The choice of processing workflow affects downstream differential methylation analysis

Depending upon the aim of a particular analysis, the absolute level of methylation might not be as important as the identification of differentially methylated loci (DMLs) or regions (DMRs) between experimental conditions or subgroups. To address this in our study, we devised a synergistic approach of assessing the workflow differential methylation impact that combines two differential methylation-centered metrics (**Figure 6a**). Based on the tumor-normal pairs included in our study, we investigated whether various workflows affected the accuracy of DMR identification with a standard DMR caller. As an external reference, we used Illumina HumanMethylation450 array data for all six tumor-normal pairs of the original study [21] to increase statistical power compared to our two sample pairs. Because the array does not cover as many CpG sites as genome-wide sequencing, we restricted our comparison to the CpG sites covered by the array. We applied the same DML identification procedure (see Materials and Methods) to all workflows and calculated AUC scores to evaluate the performance of the workflows (**Figure 6b**). As a second metric for differential analysis, we used the correlation between tumor-normal beta-value differences obtained from sequencing and microarray data (**Figure 6c**).

**Figure 6:**
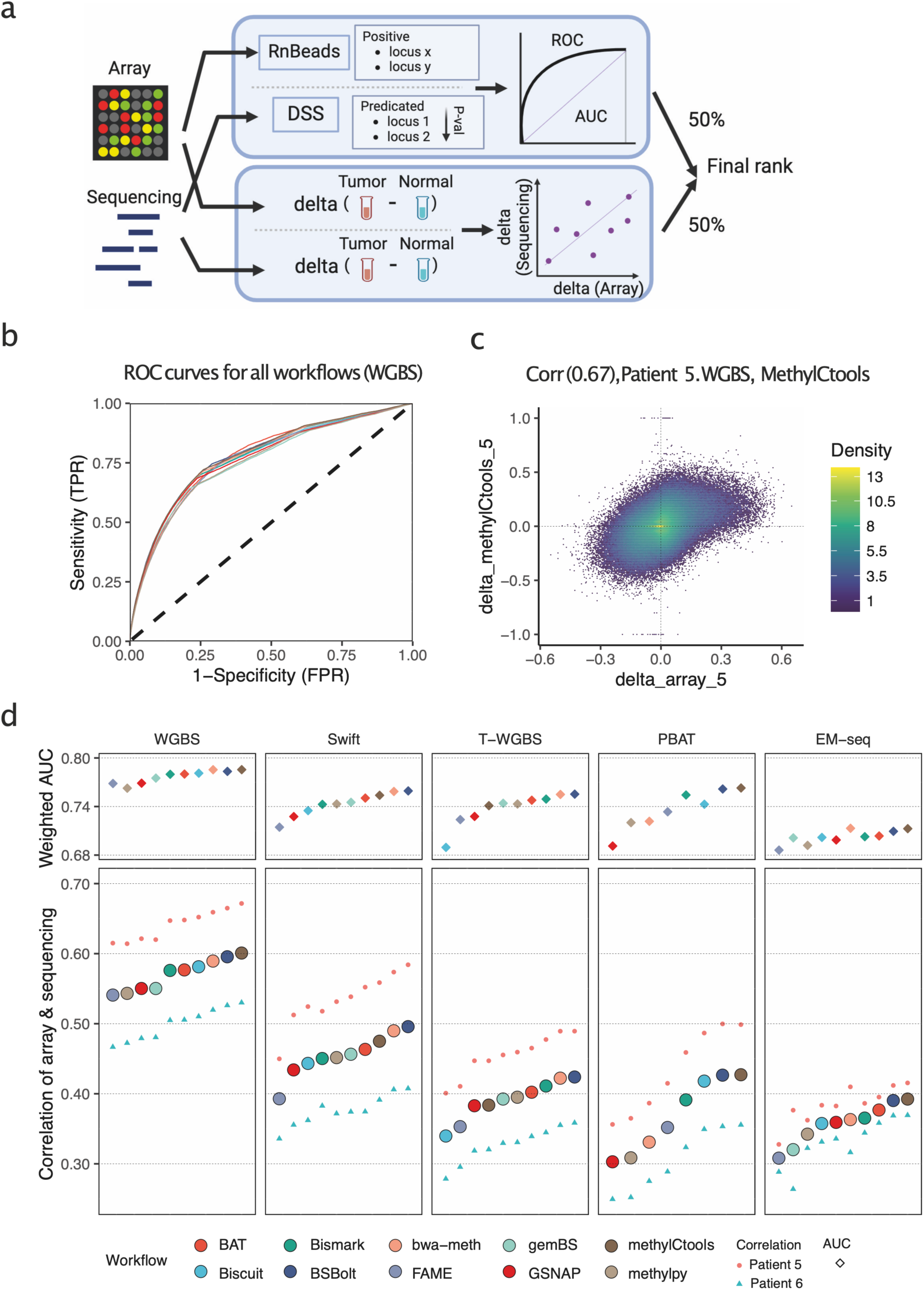
Effects of data processing on differential methylation analysis. **a.** The flowchart of the differential methylation analysis. Two types of performance measurements were employed. First, we took the differential methylation loci (DMLs) as based on Illumina HumanMethylation450 arrays for six colon cancer tumor-normal pairs as ground truth and compared them to the DMLs acquired on the sequencing data by each workflow. After applying the same filtering rules, the predicting power was estimated by ROC curves. Second, pairwise differences between tumor and normal samples were calculated both for array data, as well as for sequencing. The correlation between these differences served as the second metric. The final result was determined by averaging the ranks from two metrics. **b.** ROC curves for all workflows, based on WGBS data. Color code as in d. **c.** An example 2D density correlation plot of methylation delta betas (tumor-normal) in microarray and sequencing data for patient 5, WGBS protocol and *methylCtools* workflow. **d.** The weighted AUC and correlation of delta beta. The upper plot shows the AUC score and the lower plot shows the correlation of delta. Within each protocol, the workflows were sorted in ascending order based on their correlation values.

Regarding the AUC scores, all workflows demonstrated the capability of achieving an AUC value of at least 0.68. However, when considering correlation to Infinium array data, only in the case of WGBS the correlation coefficient surpassed 0.5, possibly indicating the increase of consistency with higher sequencing depth. Nonetheless, we noted a high degree of consistency between these two metrics across various protocols, both in terms of ranking and differences between workflows (**Figure 6d**). To integrate both assessments, we established a composite metric by combining the weighted AUC and the correlation to evaluate the effectiveness of differential methylation identification. The final DMR performance metric was determined by averaging the rankings of the workflow based on the two metrics.

Overall, we conclude that, although inferior to the effect of the sequencing depth, the workflow choice did have a measurable impact upon the differential methylation analysis. Given that the experimental protocol and depth of sequencing are often chosen *a priori* out of sample availability and sequencing resources considerations, workflow users should carefully consider their data processing strategy.

### Workflows show drastic differences in computational performance

Computational efficiency is one of the major software selection criteria in practice. To help users identify workflows that fit their available computational resources, we measured the running time and maximal memory usage of each workflow (**Figures 7a** and **7b**). All workflows implement support for parallel computations. Therefore, we supplied all the resources of the computing node when measuring the resource usage. We observed significant variation between workflows in terms of both runtime and memory requirements, regardless of the protocols. The running time varied between two extremes: 4 hours for *gemBS* and 14 days for *GSNAP*. As expected, the extremely deeply sequenced WGBS protocol had the longest running time in most workflows. Excluding WGBS, PBAT required a slightly longer running time than other protocols. We suspect that handling four different strands in PBAT leads to more computing. Overall, *gemBS*, *FAME* and *Biscuit* were the fastest workflows, ranking among the top three across all tested scenarios. Operating memory usage ranged between 16 GB for *Bismark* and 319 GB for *GSNAP*. *Bismark* had the consistently best memory footprint in all tests, followed by *FAME* and *methylpy.* Notably, the tools with perfect memory footprint oftentimes showed very large runtime and vice versa, indicating that the availability of respective resources can be decisive for the choice of workflow.

**Figure 7:**
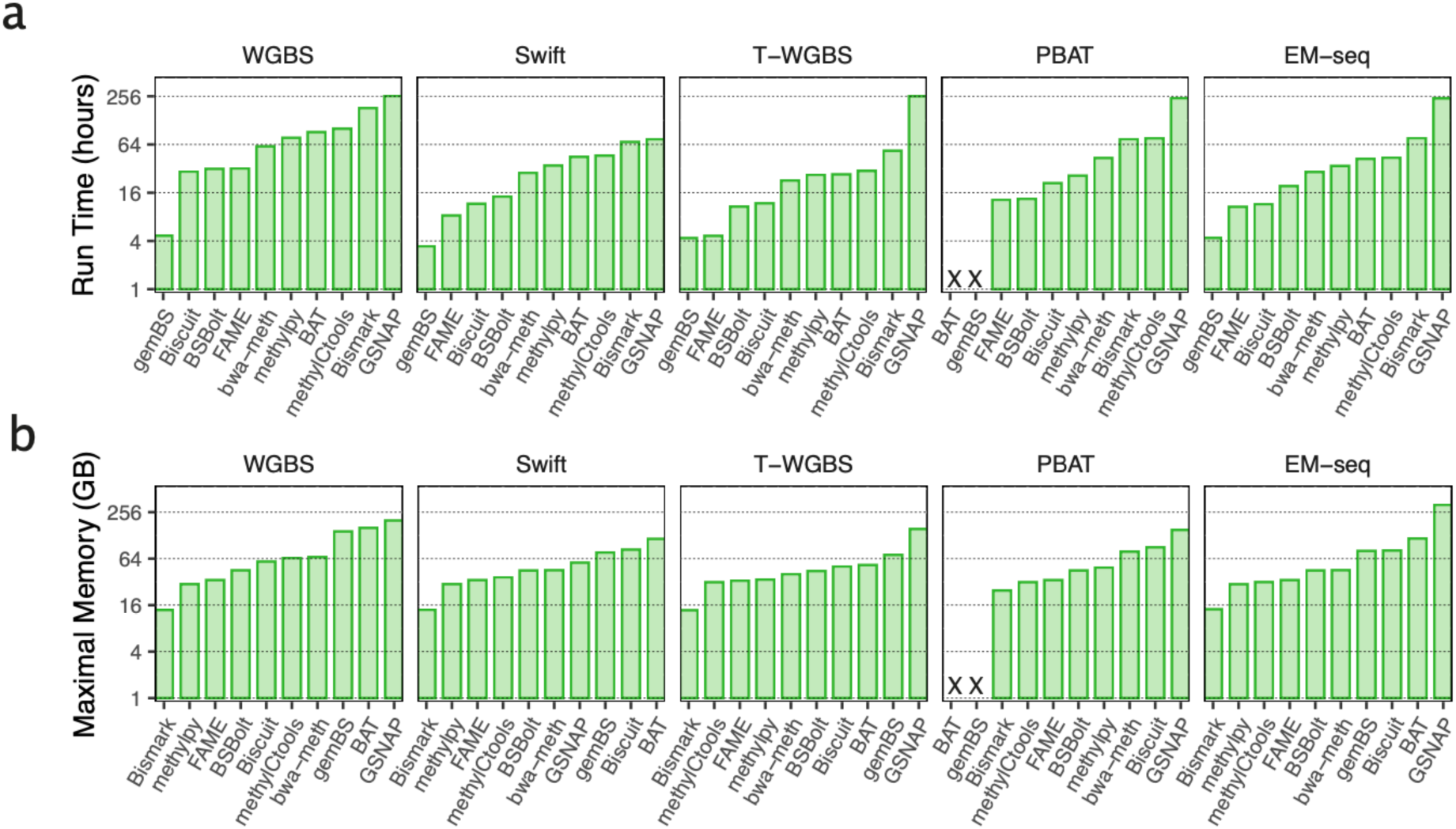
Computing resource requirements. **a.** The runtime in hours for processing sample 5N. **b.** The peak operating memory usage in gigabytes (GB) for processing sample 5N.

### Final performance ranking and recommendations

To offer guidance in selecting a bisulfite sequencing workflow, we combined the above evaluation results to create a single ranking that reflects the overall performance. We ranked the workflows based on each evaluation criterion. This ranking reflects the performance of each workflow in this specific category (see the Methods section for details). The final ranks were then calculated by averaging the individual ranks (Supplementary Table 5). Ranking was derived across all protocols considering six metrics (**Figure 8a**) as well as individually for each protocol (**Figure 8b**).

**Figure 8:**
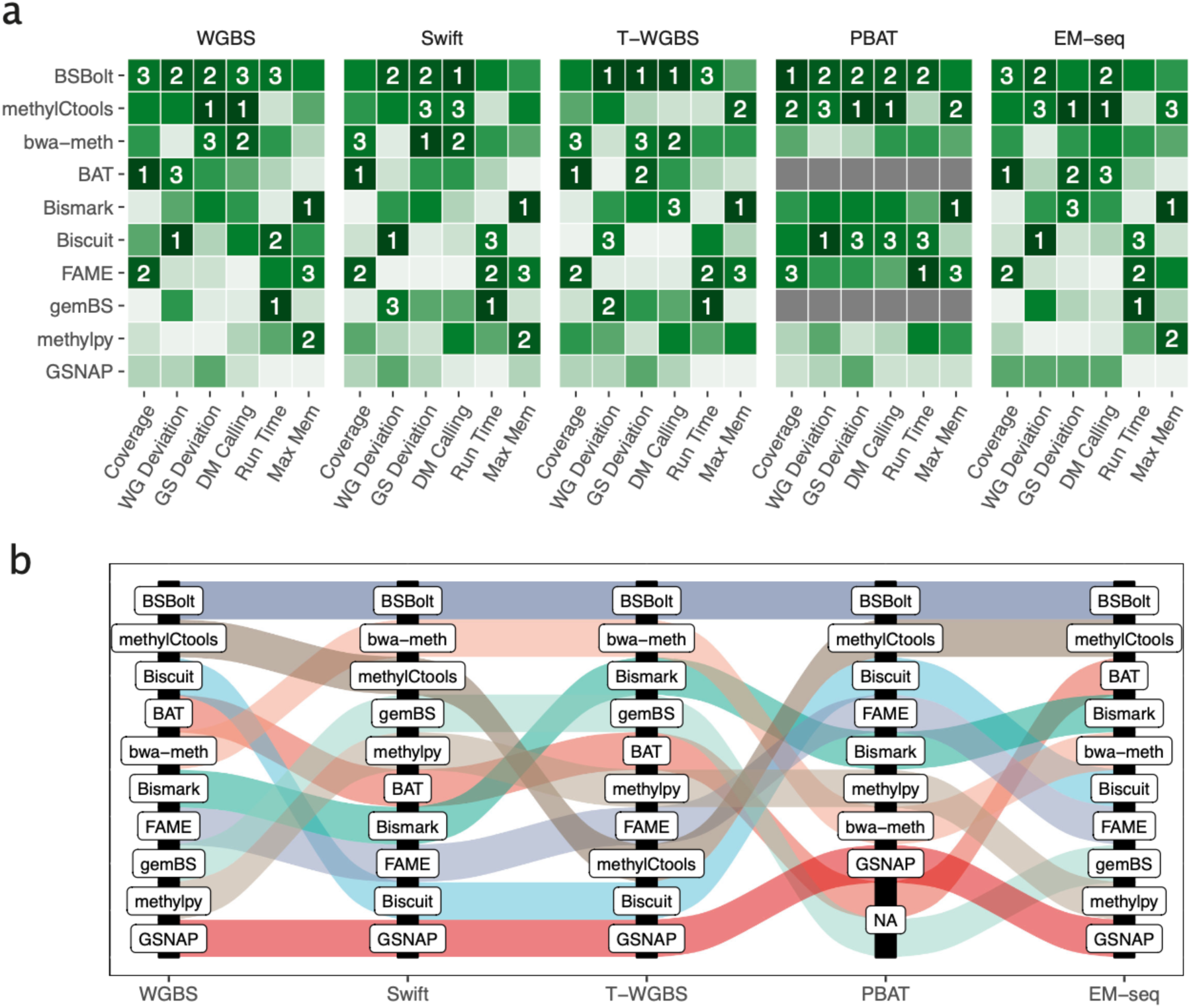
Final performance ranking. **a.** Summarized results of the benchmarking study. Color scale reflects the rank of the workflow for each metric (1 to 10, 1 is the best; dark green to light). Six metrics were incorporated, as follows. (1). ‘Coverage’: area under the curve of the ‘%CpGs covered over coverage threshold’ plot (Figure 3c). (2). ‘WG deviation’ represents the deviation across the whole genome (Figure 4c), while (3). ‘GS Deviation’: the deviation observed at the 46 gold standard loci (Figure 5c). (4). ‘DM calling’: the average of weighted AUC and the correlation of array and sequencing (Figure 6c). (5). ‘Run Time’: execution time, (6). ‘Max Mem’: the peak memory usage (Figures 7a and 7b). The workflows were ordered by the rank average of all measurements and numbers mark the first three workflows by comparison. Taking all factors into consideration, the top three performers are *BSBolt*, *methylCtools*, and *bwa-meth*. b. Alluvial plot shows the rankings for each protocol. *BSBolt* ranked first in all five protocols. Following that, *methylCtools* achieved second place in three protocols and third place in one protocol. The subsequent rankings were held by *bwa-meth*, *BAT*, *Bismark*, and *Biscuit*, which each secured a top three position in at least one of the five protocols.

According to our results, *BSBolt* ranked first among all five sequencing protocols. Following this, *methylCtools* achieved second place in three protocols and third place in one protocol. The subsequent rankings were held by *bwa-meth*, *BAT*, *Bismark,* and *Biscuit*, each of which secured the top three positions in at least one of the five protocols. The final summarized ranking considers all metrics with equal weights. This is usually not the case in real-world studies; for example, small differences in coverage depth are not important if the workflow is accurate. Therefore, we created an interactive interface that presented the results of our study and helped customize the workflow ranking. This tool is available at https://epigenomics.dkfz.de/PipelineOlympics/shiny.

### Continuous benchmarking platform for bisulfite sequencing workflows

As our final step, we aimed to turn our evaluation of 10 workflows into a continuous (“living”) benchmarking study. For this, we chose to provide a web service for workflow developers to quickly assess quality and overcome usual challenges, such as the availability of performance evaluation datasets, reliable metrics, and convenient execution environments. We implemented our continuous benchmarking platform using German public de.NBI IAAS cloud. As portal to our platform, we employed workflUX (https://github.com/workflux/workflUX), a user-friendly web service we earlier developed for running workflows implemented in Common Workflow Language (CWL). In brief, future developers can implement their workflows, or their simple single-command wrappers, in CWL and upload them into worklfUX for evaluation. Continuous benchmarking processes these uploads using downscaled versions of our benchmarking datasets and compares the results to ten existing workflows in our study, offering valuable feedback for future improvements. Usage instructions can be found on the website for reference This service is deployed on the deNBI.cloud and will be made available to developers at no cost at https://epigenomics.dkfz.de/PipelineOlympics/workflux.

## Discussion

Here, we present the results of our systematic benchmarking to provide an overview of the available complete workflows for bisulfite sequencing data and thoroughly evaluate their performance. We included regularly maintained, open-source workflows that cover different existing approaches and chose 10 to be included in our study (**Figure 1a**, Supplementary Table 1). Since different library preparation protocols have different technical and methodological aspects, we included five approaches: besides the originally developed WGBS, bisulfite-free (EM-seq) and low-input protocols (T-WGBS, SWIFT, PBAT). In addition, we present a dynamically extendable framework to include any number of additional tools, aiming to help both developers and users to evaluate their workflows of choice.

The main challenge of benchmarking studies is the lack of established ground truth measurements. Simulated data often do not capture all known and latent sources of bias and noise, whereas, for real-world data, the true underlying methylation levels are unknown. To tackle this problem, we used DNA samples from an earlier technology benchmarking study, providing highly accurate consensus DNA methylation measurements for a selection of CpG sites in tumor and normal samples. Using several genome-wide and locus-specific metrics, we objectively ranked the performance of data processing workflows in a protocol-specific manner.

Our overview of DNA methylation data generated using different protocols revealed significant protocol-specific challenges, despite the overall consistency of the generated methylation data. In particular, we identified a non-uniform coverage with a bias towards GC-rich regions in the ultra-low input PBAT protocol, causing a sensible shift in the methylation ratio. Although not in frequent use anymore, PBAT is related to modern low-input and single-cell bisulfite sequencing protocols. Our findings emphasize the need for careful workflow selection for each type of sequencing data, particularly for the low-input protocols.

The integration of all our benchmarks into the final polyathlon-style ranking showed a largely consistent workflow performance pattern across the considered experimental protocols and allowed the identification of three major performance groups. *BSBolt*, *bwa-meth,* and *methylCtools* showed reproducibly high performances for almost all criteria and protocols. Although the size of our workflow set does not allow for a rigorous statistical evaluation, it is worthwhile to note that all three best-performing tools use a reduced alphabet (3-letter) bisulfite alignment strategy with the BWA algorithm as their underlying alignment engine. In the mid-performing group, *BAT*, *Bismark*, *gemBS*, and *FAME* showed good performance, in particular with respect to runtime and memory usage, whereas *methylpy* and *GSNAP* performed worse in many metrics. *Biscuit* showed the most variable performance across experimental protocols, yet performed very well in PBAT, which is most similar to single-cell bisulfite sequencing protocols. Notably, we detected remarkable differences in CPU time and observed the trend that the workflows showing high levels of runtime and memory usage efficiency – *gemBS*, *FAME*, *Biscuit* and *Bismark* – tended to score lower in the accuracy benchmarks, revealing two diverging pathways in the evolution of workflows.

Our results show that proper workflow selection is more important with shallowly sequenced low-input protocols, since low sequencing depth amplifies the methodological and implementation-level pitfalls of each workflow. The final choice of workflow depends on individual priorities and can be affected by multiple factors. Therefore, we developed an interactive web application that visualizes the results and adapts ranking to individual importance weights.

Given the constantly accelerating influx of new bioinformatics software, comprehensive and objective benchmarking is becoming crucial. New software tools appear regularly and there are many possible combinations of tools throughout the workflow. Therefore, we introduced a dynamically expandable benchmarking platform allowing for easy addition of new tools and workflows, with the main goal of helping future developers optimize their tools. Taken together, our study helps a broad range of users to choose the proper workflows adjusted for their needs and allows expansion to further upcoming data processing tools and workflows, establishing the concept of dynamic benchmarking. This can serve as an example for benchmarking software for other data types, helping to increase the quality and reproducibility of data analysis.

## Methods

### Sample acquisition

In our study, we obtained genomic DNA isolated from two pairs of fresh-frozen colon cancer tissue samples with adjacent normal from the BLUEPRINT technology benchmarking study, Patients 5 and 6 [21]. A detailed description of the samples and the DNA isolation protocol are available in the original study.

### Library preparation and sequencing

We used five different whole-methylome sequencing protocols as outlined in Supplementary Table 2. Library preparation of T-WGBS and PBAT was performed at the Division of Cancer Epigenomics, while library preparation for WGBS, SWIFT, and EM-seq as well as the DNA sequencing were performed by the Genomics and Proteomics Core Facility at the German Cancer Research Center (DKFZ). All kits were used following the manufacturer’s instructions unless otherwise specified. It is important to note that the sequencing libraries were not prepared at the same time due to different protocol becoming available at a later time point, and this delay might have resulted in differences in sample degradation, especially affecting sample 6N in EM-seq.

#### Whole genome bisulfite sequencing (WGBS)

The amplification-free library preparation combined the EpiTest Bisulfite Kit (Qiagen) and the TruSeq DNA Sample Prep Kit (Illumina) and uses 2 μg of DNA per sample. Two lanes per sample were used on an Illumina HiSeq X Ten sequencing machine.

#### Swift Bio Accel-NGS

The Swift protocol [67, 68], as an alternative to PBAT, utilized Swift’s proprietary Adaptase instead of random priming. 200 ng DNA was used to perform bisulfite treatment followed by library preparation using the Accel-NGS-Methyl-Seq Kit (Swift Bio). For sequencing, one lane on HiSeq X Ten was used for each sample. Hereafter, we refer to this protocol as Swift.

#### Tagmentation-based whole-genome bisulfite sequencing (T-WGBS)

The details of the T-WGBS protocol have been described elsewhere [32]. Here we used an input of 30 ng DNA. Four independent libraries were constructed for each sample to reduce the impact of the PCR amplification. All libraries and samples were equally allocated and sequenced on two HiSeq2000 (Illumina) lanes.

#### Post-bisulfite adapter tagging with random priming (PBAT)

PBAT libraries were prepared as previously described [69] using a customized protocol for ultralow input materials based on the single-cell bisulfite sequencing protocol [70]. In brief, 6 ng of purified DNA was subjected to bisulfite conversion, a single pre-amplification for 90 min at 37°C, adaptor tagging, and finally 14 cycles of library preparation. Libraries were purified using double 0.7 × SPRI selection and sequenced on a HiSeq X Ten sequencer, applying 150 base pairs (bp) paired-end sequencing at the DKFZ Genomics and Proteomic Core Facility in Heidelberg.

#### NEBNext Enzymatic Methyl-seq based on TET and APOBEC

Bisulfite-free library preparation was performed with the NEBNext Enzymatic Methyl-seq (EM-seq) Kit [71] using 50 ng of DNA. Each sample was sequenced using one lane on a HiSeq X Ten.

The detailed information about read sets generated in this study including the list of FASTQ files, along with the total number of sequences and bases, read lengths, mean PHRED scores, non-CpG methylation levels, conversion rates, as measured by *Bismark* are given in Supplementary Table 2.

### Workflow selection and deployment

The methylome analysis workflows were selected after a careful review of the literature and publicly available software (Supplementary Table 1). The main criteria were as follows: the latest update time was after 2020 and the number of citations per year was at least 10. In addition to the qualifying list, we added two recent workflows, *Biscuit* and *FAME*, and one well-established workflow frequently used by collaborators (*BAT*). The final selected workflows are the following: *BAT* [60], *Biscuit* [61, 72], *Bismark* [40], *BSBolt* [62]*, bwa*-*meth* [51], *FAME* [49], *gemBS* [53], *GSNAP* [63], *methylCtools* [52], and *methylpy* [64]. Containerization and the CWL workflow language were used to enhance stability and reusability. Each component of the workflow was implemented as a Docker container and the pipeline was implemented in a CWL wrapper. The CWL ran on a standardized virtual machine (VM) equipped with the following specifications: Centos 7.9, 2 x 14-core Intel Xeon E5-2660 v4 (56 threads), and 512 GB RAM. The processing times and maximum memory requirements were collected from the job notification reports of the IBM Spectrum LSF platform.

### Protocol-specific differences in the processing workflows

All computational workflows consisted of the following steps: read preprocessing, alignment, post-processing, and methylation calling. A detailed description of the workflows, version numbers, and individual steps, the protocol-specific parameter settings adapted for each protocol are described in the following sections, with detailed information listed in Supplementary Table 6.

#### Trimming

Following the manufacturer’s recommendation, the random 3’-tails added by the Adaptase during the Swift protocol were removed. The removed segments were the last 15bp of the R1 and the first 15bp of the R2. In T-WGBS, the Tn5 transposase creates a short 9bp gap at the 3’ end cutting site, which is subsequently repaired by DNA polymerase and DNA ligase. During this repair process, nine base pairs are incorporated at the end of the R2 using unmethylated cytosines. Therefore, these base pairs should not be used for methylation calling but can be used for alignment. The unmethylated regions were not considered during methylation calling in the workflows supporting this, like *Bismark*, *bwa-meth*, *GSNAP*, and *methylCtools*. For the others, additional hard trimming was applied to remove the base pairs before alignment.

PBAT is a non-directional protocol that requires special handling. *BAT* works only with directional protocols; therefore, it was not run with PBAT. Although *gemBS* supports non-directional protocols, we were faced with an error that we were unable to resolve with the help of the authors until the manuscript was written. Thus, *gemBS* was not run on PBAT data. During preprocessing, random hexamers (first and last 6 bps of R1 and R2), which were added during the two steps of random priming, were removed. *MethylCools* provides a patch script to support PBAT, which determines the strands of read pairs R1 and R2 obtained from a nondirectional protocol (available from https://github.com/cimbusch/TWGBS). We ran this script on PBAT raw data to create the input for *methylCtools*.

#### Alignment

Bisulfite treatment of DNA, followed by PCR amplification, can produce four (bisulfite-converted) strands for a given locus. Depending on the adapter used, two different approaches were employed for the construction of BS-Seq libraries. In directional bisulfite sequencing, the library construction process selectively sequences either the original top strand (OT) or the original bottom strand (OB) of the DNA. Consequently, only reads aligned to OT and OB are considered, whereas alignments from the complementary strands (CTOT and CTOB) are typically not generated. In contrast, nondirectional bisulfite sequencing includes all four strands generated in the bisulfite-converted PCR process (OT, CTOT, OB, and CTOB) in the sequencing library with roughly equal likelihood, which means that alignments to all four strands are considered valid. When executing the analysis for PBAT, we configured the directional sequencing parameters for the four protocols using non-directional sequencing.

#### Removal of PCR duplicates

All workflows, except *BAT* included a duplicate removal step. The principle is to perform duplicate removal on the libraries independently. For WGBS, the two lanes sequencing the same library were aligned separately, merged, and deduplicated. For T-WGBS, which featured four independent libraries, the libraries were aligned and deduplicated individually.

#### Methylation calling

As mentioned in the trimming section, some workflows have the possibility to ignore bases at the end of the reads during methylation calling. Apart from the case specifically mentioned for T-WGBS, other protocols might also benefit from such an approach. Ideally, we expect the probability of observing a methylated C to be constant across any given read. However, an increase or decrease in methylation level is often observed, especially at the end or at the beginning of the reads. This methylation bias or M-bias can lead to an overestimation of DNA methylation. Therefore, during methylation calling, the methylation status of the last several base pairs of the reads should be ignored. The exact length of the ignored region depends on the protocol and other technical parameters. Two workflows provide a function to determine the sequence affected by M-bias. The caller of *bwa-meth*, *MethylDackel* provides an automatic M-bias identification tool (*MethylDackel* mbias), whereas *Bismark* provides a function (coverage2cytosine) with R scripts to generate M-bias plots. Therefore, we applied sample-specific alignment trimming for these two workflows. The exact number of base pairs subjected to alignment trimming is listed in Supplementary Table 6. Other methylation callers do not specifically address the M-bias.

### Statistical testing

In the genome-wide deviation analysis, we employed a paired t-test to determine if the beta values of the protocol exhibited consistent underestimation tendencies. Similarly, for the deviation of preselected loci, we conducted a two-way Analysis of Variance (ANOVA) to assess the joint impact of samples and workflow procedures on accuracy. In the context of assessing the correlation between alignment rate and whole-genome deviation in PBAT, we utilize the ‘corr.test’ package in R to calculate the Spearman correlation and test for its significance.

### Benchmarking metrics

Methylation calls in .bed or .bedgraph format were imported using the R package *methrix* [66], giving *BSgenome.Hsapiens.UCSC.hg38* as a reference. All downstream analyses were performed on the summarized methylation and coverage matrices using R 4.1.

#### The area under the depth-vs-coverage dependency curves

The area under the curve was calculated based on the dependency of the coverage fraction at a given cutoff (**Figure 3c**), where the x-axis is the read coverage threshold, and the y-axis is the cumulative fraction of covered genomic CpGs. The AUC score only considers read coverage from 0 to 200 and is normalized to 1 by dividing it by 200.

#### Genome-wide deviation from data-driven consensus

We utilized a data-driven approach to create a consensus corridor for all the CpGs. This was achieved by employing three high-coverage protocols, including WGBS, Swift, and EM-seq. The consensus corridor was defined as the smallest region encompassing at least five measurements from each protocol. The mean absolute deviation from the pre-created reference among all CpGs determines the ranking of workflows (**Figure 4d**).

#### Deviation from the consensus corridor of 46 preselected loci

To establish consensus methylation calls without the need of using simulated data we preselected loci from a previous benchmarking study conducted by the Blueprint Consortium [21]. In this study, 48 regions (16 mandatory and 32 recommended) were selected based on genome-wide methylation screen and each selected region was analyzed using multiple technologies by different labs. We excluded two regions (recommended 29 and 30) due to a lack of data points (less than or equal to three) leaving 46 regions, summarized in Supplementary Table 4. Following the original approach, we identified the consensus corridor as the narrowest interval, with measurements from three different technologies, adding a 5% flanking region. The absolute deviation was calculated for each method, workflow, and sample, based on each region. The difference *d_w_i_s_j_r_k__* was calculated as follows:

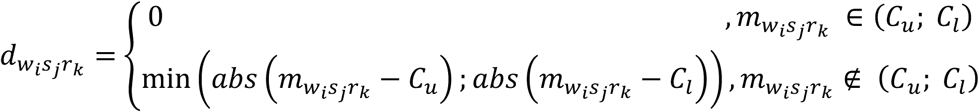

where *m_w_i_s_j_r_k__* is the methylation level of the workflow i and sample j at the region k, *C_u_* and *C_l_* are the upper and the lower border of the consensus corridor. The mean absolute deviation from the consensus corridor of each protocol-workflow pair was then calculated as 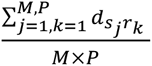, where *M* and *P* are the number of samples, and regions (**Figure 5c**).

#### Differential methylation analysis

Combined metrics were established to assess the accuracy of differential methylation detection. First, it was treated as a classification problem and evaluated using the standard weighted area under the curve metric. The targeted methylation assay benchmarking study conducted by the BLUEPRINT Consortium also made the data from Illumina Human Methylation Arrays available. This dataset comprised six pairs of colon tumor-normal samples. Two of these pairs were included in our study; therefore, we used these data to estimate the accuracy of differential methylation calling on the sequencing-based results. Differential analysis of the arrays was performed using *limma* [73], as implemented in the *RnBeads* R package [74], in a paired setting. We used *DSS* [75] to analyze differential methylation of the sequencing-based data in a non-paired setting because DSS required at least three samples for the paired mode. In both differential lists, a false discovery rate (FDR) of 0.1 was used as the threshold to determine significant differences. The AUC score for the hypermethylation and hypomethylation events was calculated separately and combined into a weighted score with the number of events (**Figure 6b**). The second part of the metrics is the Pearson correlation of delta beta values (beta value difference between normal and tumor samples) from the array and sequencing. The beta value of the microarray was extracted using RnBeads, and the correlation between the delta values of the two patients was used as a metric (**Figure 6c**).

#### Compute runtime and memory usage

The DKFZ Cluster Center allocated a few exclusive computing nodes for the project to ensure an identical and isolated environment. The computing nodes were equipped with 56 CPUs and 256 GB RAM. To estimate the CPU time and maximal RAM usage, we ran all protocol-workflow pairs on a normal sample of Patient 5 in this environment. The tasks were submitted to the IBM Spectrum LSF (Load Sharing Facility) platform. The values of “Run Time” and “Max Memory” in the job notification report were used as the metrics. Unfortunately, *Bismark* ran prohibitively slowly in this cluster environment with distributed network storage compared to single-node mode; therefore, we estimated the values through a down-sampling approach, including six subsamples ranging from 5% to 30%, with increments of 5% each. We processed these subsamples using *Bismark* and, based on the results, applied linear regression to estimate the overall execution time and memory usage.

### Ranking

The rank average of all the measurements was used to summarize the results of the benchmarking study. For each metric, the rank scale ranged from 1 to 10, with 1 indicating the best. Each workflow has a rank score calculated by averaging all measurements across the five protocols.

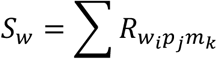

The rank average is calculated as follows, where the *R_w_i_p_j_m_k__* is the rank of workflow i in protocol j, and metric k. If multiple workflows have the same rank average, then the average z-score for each metric is used as the secondary ranking basis.

### Shiny app details

Shiny is an R package that simplifies the creation of interactive web applications and dashboards directly from R code. It is seamlessly integrated with R, enabling real-time interactivity and responsive data-driven applications. We utilized Shiny to construct our rich data website to share our research findings, allowing users to customize how they access results according to their interests. We provided various statistics, such as methylation and coverage, along with detailed visualizations of 46 gold-standard loci and a customizable ranking table. The shiny application is available at https://epigenomics.dkfz.de/PipelineOlympics/shiny/

### WorkflUX server details

workflUX, formerly known as CWLab, is an open-source web application designed to streamline the deployment of big data workflows. Its standout features include platform versatility and seamlessly operating on Linux, MacOS, and Windows, ensuring compatibility with the preferred operating system. Furthermore, it offers support for containerization, including Docker, singularity, and udocker, enabling efficient dependency management and simplifying the workflow deployment process. workflUX seamlessly integrates with a range of CWL runners, such as cwltool, Toil, Cromwell, Reana, and CWLEXEC, empowering users to execute CWL workflows across various infrastructures, from single workstations to HPC clusters and cloud platforms. We implemented automated benchmarking using workflUX. Workflow developers are required to implement their workflows using Common Workflow Language (CWL). The workflUX server utilizes preprepared small datasets and related CWL job configurations to create an automated benchmarking service freely available at https://epigenomics.dkfz.de/PipelineOlympics/workflux/.

## Availability of data and code

We have uploaded the raw sequencing data to European Genome-Phenome Archive under the accession EGAS50000000541.

The CWL versions of all evaluated workflows with containerization details, as well as all analysis R scripts, are publicly available at https://github.com/CompEpigen/PipelineOlympics.

## Ethical Compliance

All procedures performed in studies involving human participants were following the ethical standards of the institutional and/or national research committee and with the 1964 Helsinki Declaration and its later amendments or comparable ethical standards.

## Author Contributions

P.L., R.T., Y.A., C.P. and C.B. initiated the study and acquired funding. Y.L. implemented of all workflows on local compute infrastructure, performed data processing, analysis, interpretation, created all figures and tables and wrote the manuscript with P.L and R.T. M.E. and C.B provided samples for the study. D.W., O.M., M.Schoe., M.H., F.P. performed sequencing experiments supervised by S.W., C.G., D.B.L and C.P. L.W., G.G., F.T. and I.B. managed sequencing data and computing infrastructure. K.B., A.W., K.N., S.K., P.K., J.F., E.A.H., H.K., S.H., A.M., M.Schu. supported workflow implementation and bioinformatic analyses. Y.L., P.Lafr., M.C. developed web-services supported by S.T. and C.L. C.P., C.B., Y.A., J.W., C.G., M.H., D.B.L, V.H., M.Z., M.I., E.A.G., S.H., M.Schl., M.Schu. supported data interpretation. P.L. and R.T. jointly supervised the study. All authors read, edited and approved the final text of the manuscript.

## Acknowledgements

We are grateful to Ingrid Scholz for her help with data import, to Matthias Bieg and Charles Imbusch for sharing the implementation of the methylCtools workflow for PBAT data.

## Supplementary Figure Legends

**Supplementary Figure 1**

Schematic overview of protocol chemistries for whole-genome bisulfite sequencing (WGBS), Swift Bio’s Accel-NGS (Swift), Tagmentation-based whole-genome bisulfite sequencing (T-WGBS), PBAT with random priming (PBAT), and NEB’s NEBNext (EM-seq). The amount of DNA used in this study is specified next to each protocol name. These protocols can be categorized into three groups based on the input DNA amount, which is indicated beneath the protocol name. The color coding refers to native DNA (dark blue), cytosine-converted DNA (light blue), adaptors (orange and green) and unmethylated artificial ends (purple).

**Supplementary Figure 2**

Characteristics of methylation data obtained with different protocols. a. An extension of Figure 2a. The histogram displays the distribution of depth for the different protocols by samples. The boxplot at the bottom of the histogram indicates the median and interquartile range (IQR) values, offering insight into the central tendency and spread of the distribution. b. An extension of Figure 2b. The density distribution of the GC content of the reads by samples. Note that the read sequence itself was not used, but rather the reference genome sequence at the corresponding alignment position. c. M-bias plots show average methylation levels by read position for the samples named 5N, 6N (normal), and 5T, 6T (tumor), separately for read 1 and read 2. In the ideal case, the level of methylation is independent of the position. The read lengths vary between protocols due to differences in both trimming and the original read lengths.

**Supplementary Figure 3**

Fractions of retained reads after the trimming, alignment, and duplicate removal steps. Workflow steps are given on the horizontal dimension. All fractions are calculated based on the read number of raw data. UNK indicates the workflow does not provide the corresponding BAM files. NA indicates the workflow failed on the dataset (BAT does not support PBAT and *gemBS* failed on PBAT) or does not include this step (BAT does not contain a deduplication step).

**Supplementary Figure 4**

Distribution of methylation values in WGBS data after downsampling to PBAT coverage (extension of Figure 3c). An accentuated mode at approximately 0.8 was observed in the beta distribution of WGBS. Unlike the other four protocols, which all display two peaks at 0 and 1, WGBS does not exhibit a distinct peak at 1. This phenomenon is suspected to be a result of the higher read coverage associated with WGBS, causing the peaks to shift from 1 towards 0.8. To verify our hypothesis, we downsized the WGBS sample to 8x read coverage (matching the depth of PBAT in the study), and the down-sampled reveals a peak at position 1, with a distribution closely resembling that of the other four protocols.

**Supplementary Figure 5**

Abundance of chimeric reads in PBAT libraries. We confirmed the presence of chimeric reads, as shown in previous studies [66], which contain sequences from two or more distinct genomic loci. To assess the occurrence of chimeric reads in different protocols, we define reads that map to different chromosomes as chimeric reads and calculate their proportion relative to the total mapped reads. This analysis reveals that the proportion of chimeric reads generated in PBAT is approximately 6.64 times that of Swift and 10.33 times that of WGBS. The Y-axis represents the fraction of chimeric reads among mapped reads. The counts for mapped reads and chimeric reads are extracted from the *samtools flagstat* report. In Line 5, ‘mapped’ is used to denote the number of mapped reads, and in Line 12, ‘with mate mapped to a different chr’ is used to represent the number of chimeric reads.

**Supplementary Figure 6**

Breadth vs depth of coverage diagrams (an extension of Figure 3a). These plots display the percentage of covered CpGs (y-axis) below varying read coverage thresholds in log10 scale (x-axis). It provides a practical method for determining the proper coverage cut-off threshold.

**Supplementary Figure 7**

The distribution of beta values for sample-protocol pairs. The beta value of each workflow is represented in a separate line on each plot.

**Supplementary Figure 8**

Deviation stratified by classes of genomic regions. Hg38 annotation of R package *annotatr* (v1.14.0) was used. The mean of the deviation among all workflows sorts the annotations (x-axis). Our conclusion is that genic regions (genes_5UTRs, genes_exons, etc.) exhibit lower discrepancies values when compared to intergenic regions (genes_intergenic). The order of y-axis is the final ranking.

**Supplementary Figure 9**

Relationship between alignment rate and absolute means deviation on whole-genome scale in the PBAT dataset. Correlation suggests that a higher alignment rate is associated with greater accuracy, as indicated by a lower absolute mean deviation. The Spearman correlation and p-value on the plot represent the average of sample-wise calculations.

**Supplementary Figure 10**

Deviation plot of 46 loci (WGBS) for assessing gold standard-based methylation call accuracy. The plot displays the deviation from the gold-standard consensus corridors for these 46 selected loci. The grey boxes represent the consensus corridors, as reported by the BLUEPRINT Consortium. Each dot on the plot represents a measured beta value for a specific workflow, and the lines connecting the dots depict their respective deviations from the consensus corridors. The loci are sorted by the mean of the consensus corridor. This plot serves as an extension of Figure 5a, which only indicates 6 loci.

**Supplementary Figure 11**

Same as Supplementary Figure 10 but using down-sampled WGBS alignments to a coverage level equivalent to PBAT (8x coverage) to assess the effect of read coverage. We excluded workflows that were not run on the PBAT dataset and those that did not provide BAM files. As a result, this plot involves only 6 workflows. It represents the deviation of 46 loci in the down-sampled WGBS dataset, which now matches the coverage of PBAT.

**Supplementary Figure 12**

Same as Supplementary Figure 10, but for the Swift protocol.

**Supplementary Figure 13**

Same as Supplementary Figure 10, but for the T-WGBS protocol.

**Supplementary Figure 14**

Same as Supplementary Figure 10, but for the PBAT protocol.

**Supplementary Figure 15**

Same as Supplementary Figure 10, but for the EM-seq protocol.

**Supplementary Figure 16**

Deviations from gold standard consensus corridors (an extension of Figure 5b). The deviation from the consensus corridors (CC) of the four protocols for all samples combined, while the WGBS data is shown in Figure 5b. The pie charts on the left illustrate the proportion of data points falling outside versus inside the CC. On the right, the box plot displays the distribution of deviations, excluding data points within the CC. The columns labeled ‘N/T’ indicate whether the data pertains to normal or tumor samples, respectively. Positive values indicate prediction above the CC, while negative values indicate predictions below the CC.

**Supplementary Figure 17**

Example of a software bug resulting in erroneous methylation calls. In the case of the normal sample from WGBS patient 5, locus r23, *Methylpy* detects lower methylation beta values compared to other workflows. From (a), it can be observed that *Methylpy* determines lower beta values, and upon alignment, it is evident that both workflows have the same number of aligned reads. In (b), we observed that the intermediate BAM sequence and final sequence from *Methylpy* (3rd and 4th row) do not match raw read (1st) and BSBolt (2nd row). *Methylpy* employs a three-letter method that carries out in silico conversion of reads before alignment. In the methylation calling step, the converted reads must be restored to their original sequences. These observations are suggesting of an implementation error in the simulated bisulfite conversion.

## Supplementary Table Legends

**Supplementary Table 1**

Overview of published bisulfite sequencing processing workflows. Data including code repository URLs and publication DOIs, publication years, citation counts and other impact metrics, maintenance status etc.

**Supplementary Table 2**

Overview of five sequencing protocols used in the study and basic statistics of the generated methylation sequencing data, including PHRED score, percentage of methylated non-CpG cytosines, conversion rate and read length. Sequences of artificial adapter sequences, based on the characteristics of the protocols which were removed before the processing are given for each protocol.

**Supplementary Table 3**

Raw dataset overview. The table summarizes the information on the generated FASTQ files for each sequencing library, basic processing statistics of each workflow execution, including the fraction of CpGs covered, depth and percentage of CpG methylation.

**Supplementary Table 4**

Overview of reference loci used for the experimental gold standard evaluation.

**Supplementary Table 6**

Detailed information of the evaluated workflows and execution parameters used for each protocol.

