## Supplementary Figure for "Pipeline Olympics: continuable benchmarking of computational workflows for DNA methylation sequencing data against an experimental gold standard"

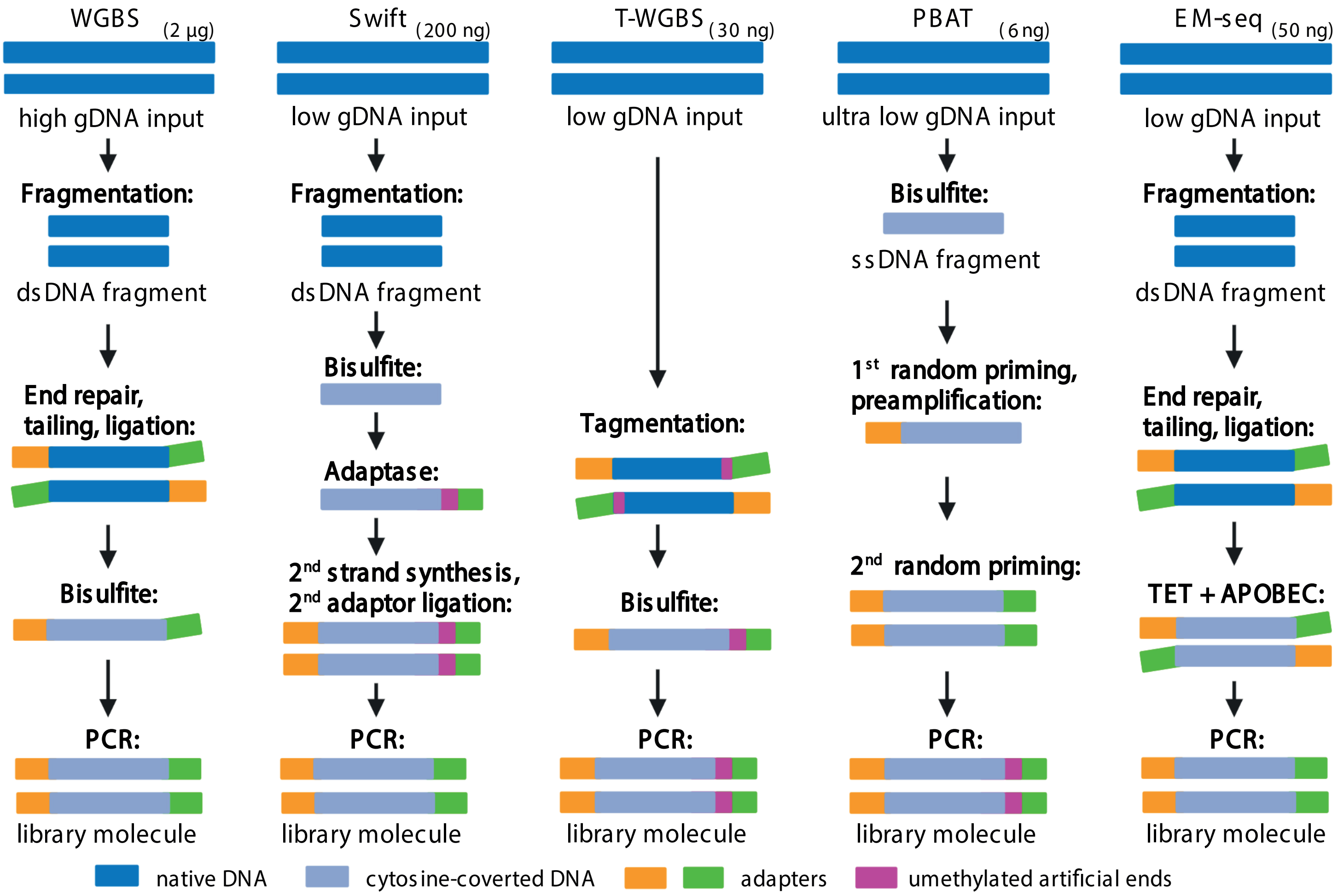


**Supplementary Figure 1:**

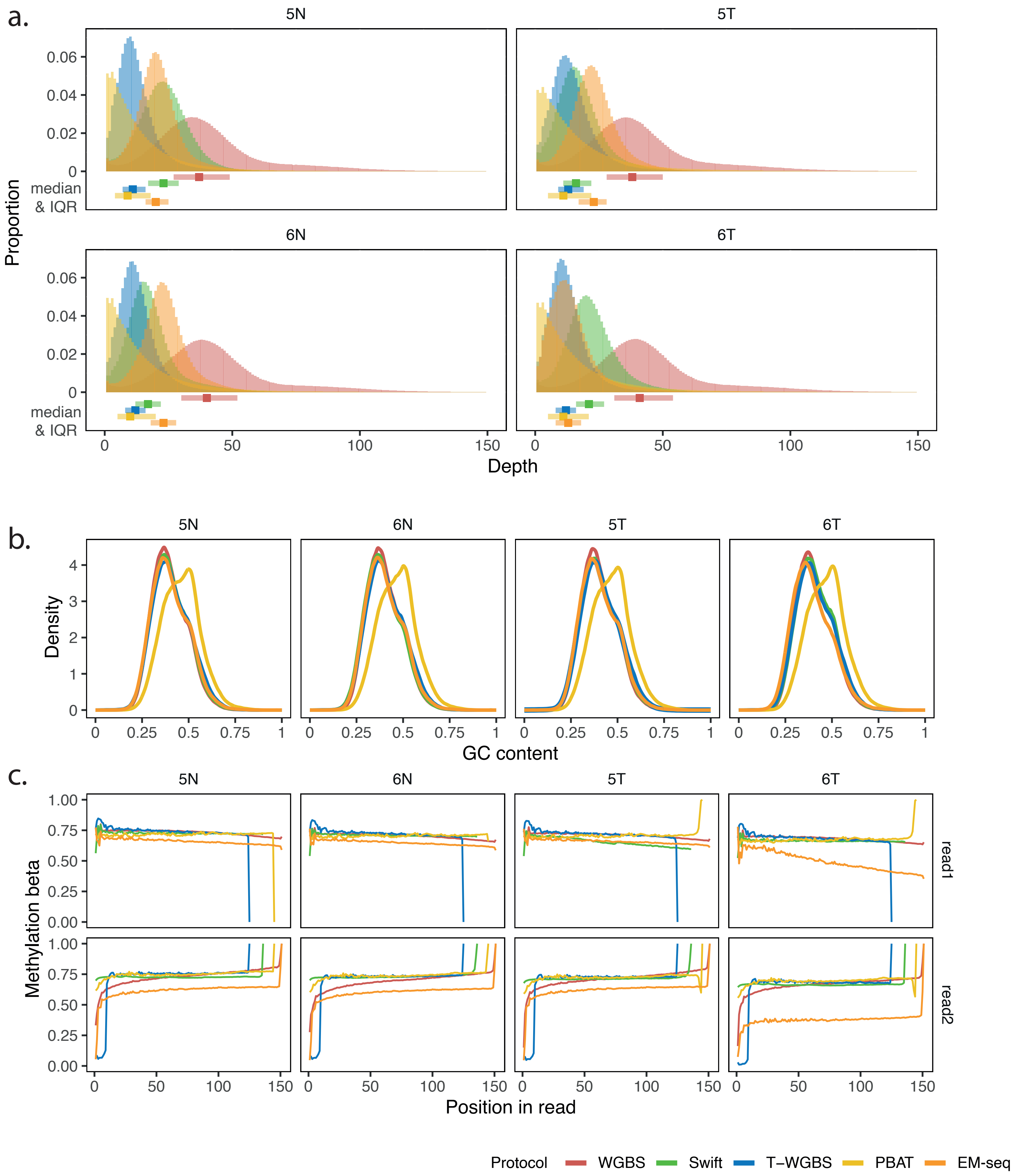


**Supplementary Figure 2:**

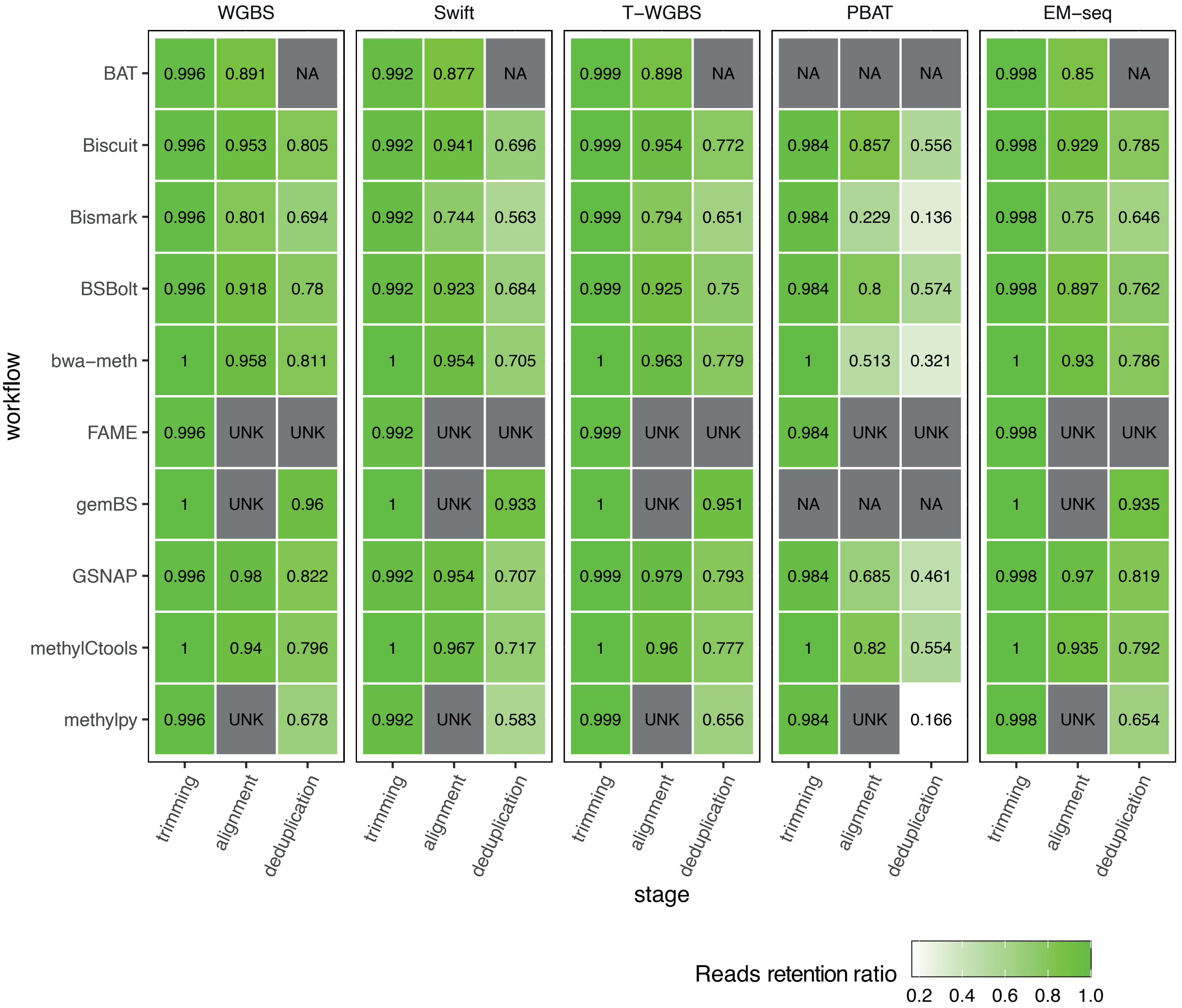


| workflow | WGBS trimming | WGBS alignment | WGBS deduplication | Swift trimming | Swift alignment | Swift deduplication | T−WGBS trimming | T−WGBS alignment | T−WGBS deduplication | PBAT trimming | PBAT alignment | PBAT deduplication | EM−seq trimming | EM−seq alignment | EM−seq deduplication |
|---|---|---|---|---|---|---|---|---|---|---|---|---|---|---|---|
| BAT | 0.996 | 0.891 | NA | 0.992 | 0.877 | NA | 0.999 | 0.898 | NA | NA | NA | NA | 0.998 | 0.85 | NA |
| Biscuit | 0.996 | 0.953 | 0.805 | 0.992 | 0.941 | 0.696 | 0.999 | 0.954 | 0.772 | 0.984 | 0.857 | 0.556 | 0.998 | 0.929 | 0.785 |
| Bismark | 0.996 | 0.801 | 0.694 | 0.992 | 0.744 | 0.563 | 0.999 | 0.794 | 0.651 | 0.984 | 0.229 | 0.136 | 0.998 | 0.75 | 0.646 |
| BSBolt | 0.996 | 0.918 | 0.78 | 0.992 | 0.923 | 0.684 | 0.999 | 0.925 | 0.75 | 0.984 | 0.8 | 0.574 | 0.998 | 0.897 | 0.762 |
| bwa−meth | 1 | 0.958 | 0.811 | 1 | 0.954 | 0.705 | 1 | 0.963 | 0.779 | 1 | 0.513 | 0.321 | 1 | 0.93 | 0.786 |
| FAME | 0.996 | UNK | UNK | 0.992 | UNK | UNK | 0.999 | UNK | UNK | 0.984 | UNK | UNK | 0.998 | UNK | UNK |
| gemBS | 1 | UNK | 0.96 | 1 | UNK | 0.933 | 1 | UNK | 0.951 | NA | NA | NA | 1 | UNK | 0.935 |
| GSNAP | 0.996 | 0.98 | 0.822 | 0.992 | 0.954 | 0.707 | 0.999 | 0.979 | 0.793 | 0.984 | 0.685 | 0.461 | 0.998 | 0.97 | 0.819 |
| methylCtools | 1 | 0.94 | 0.796 | 1 | 0.967 | 0.717 | 1 | 0.96 | 0.777 | 1 | 0.82 | 0.554 | 1 | 0.935 | 0.792 |
| methylpy | 0.996 | UNK | 0.678 | 0.992 | UNK | 0.583 | 0.999 | UNK | 0.656 | 0.984 | UNK | 0.166 | 0.998 | UNK | 0.654 |

**Supplementary Figure 3:**

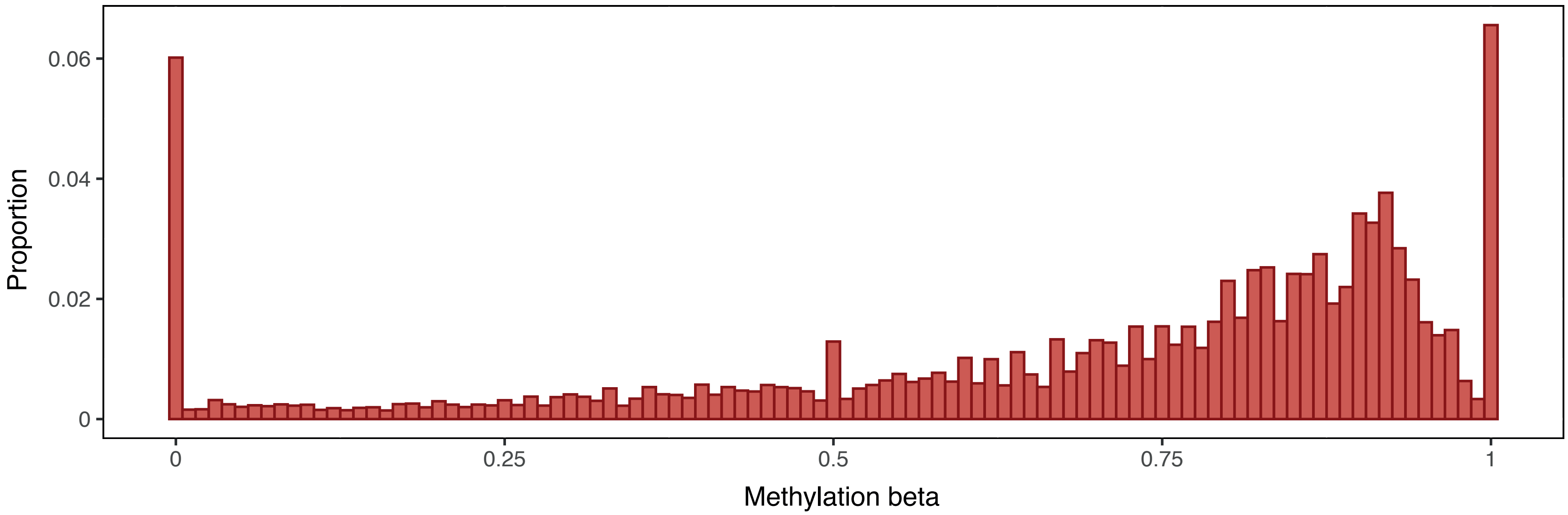


**Supplementary Figure 4:**

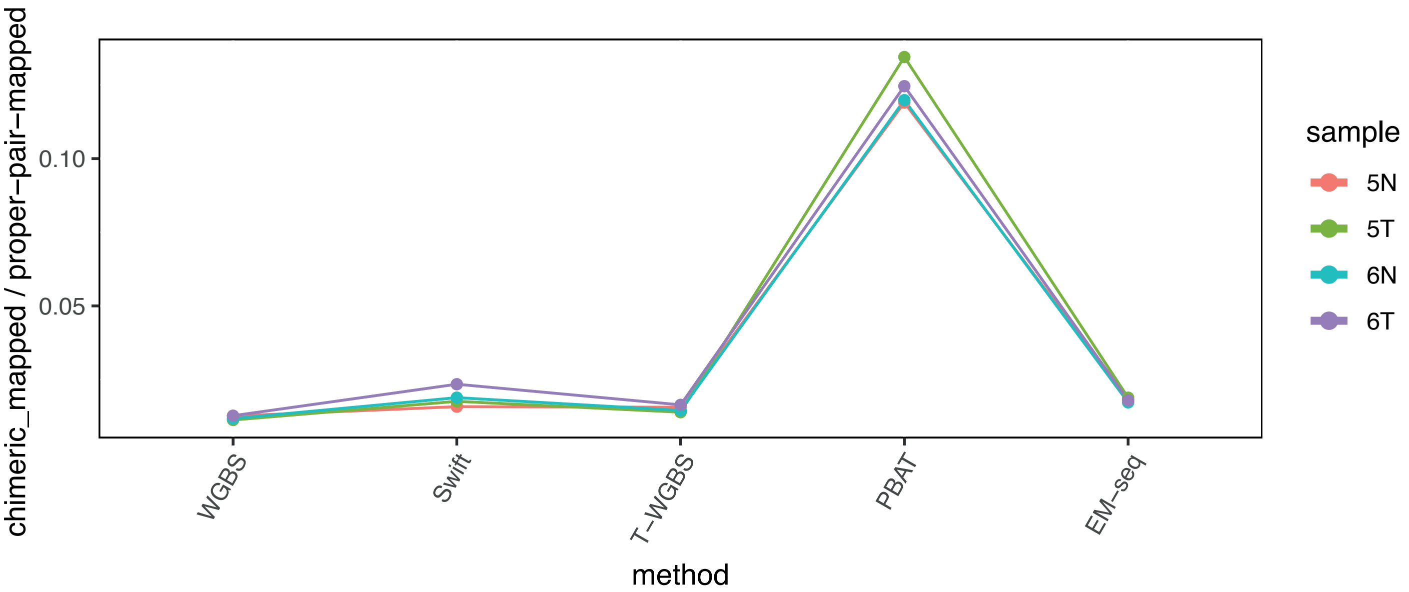


**Supplementary Figure 5:**

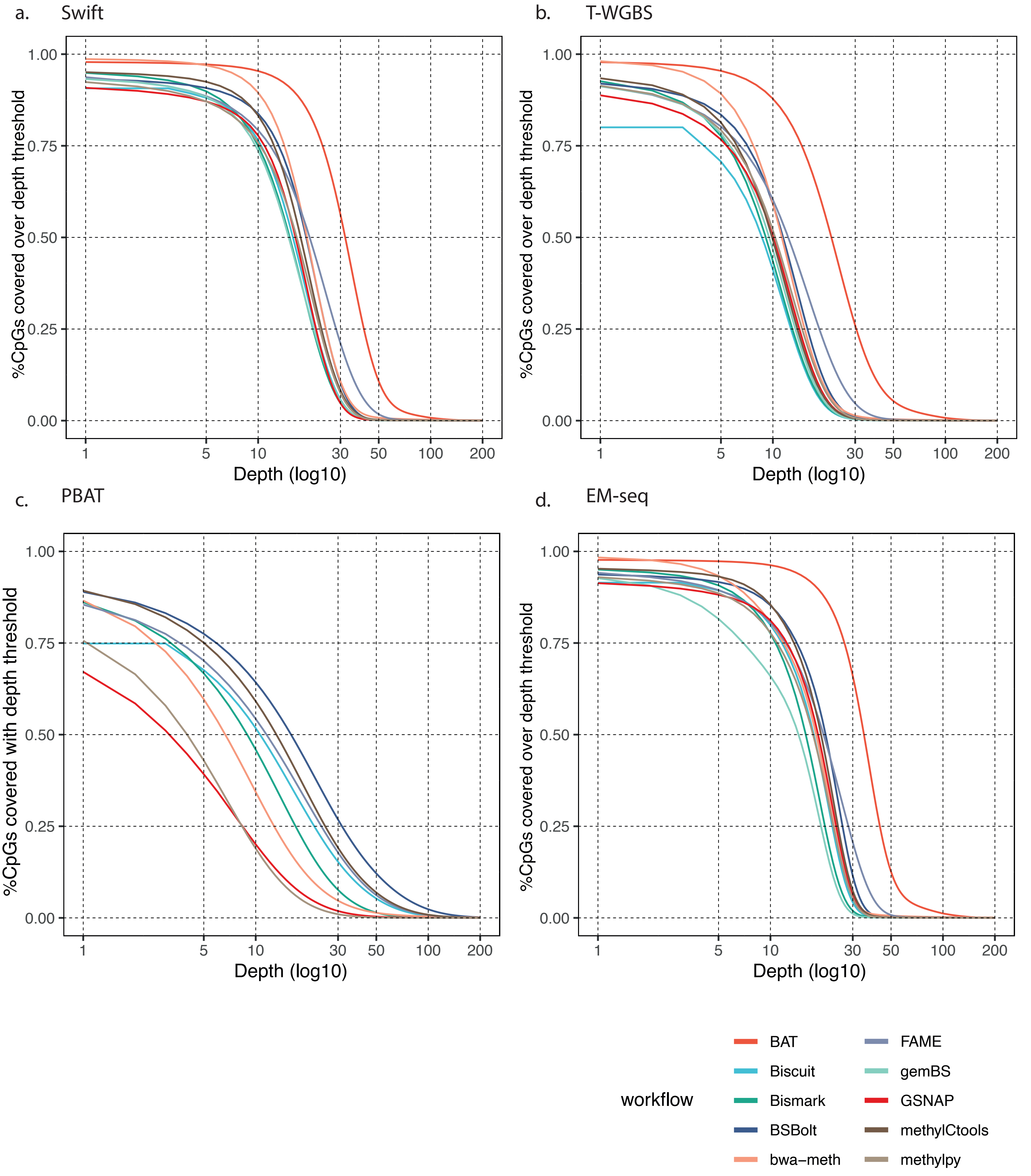


**Supplementary Figure 6:**

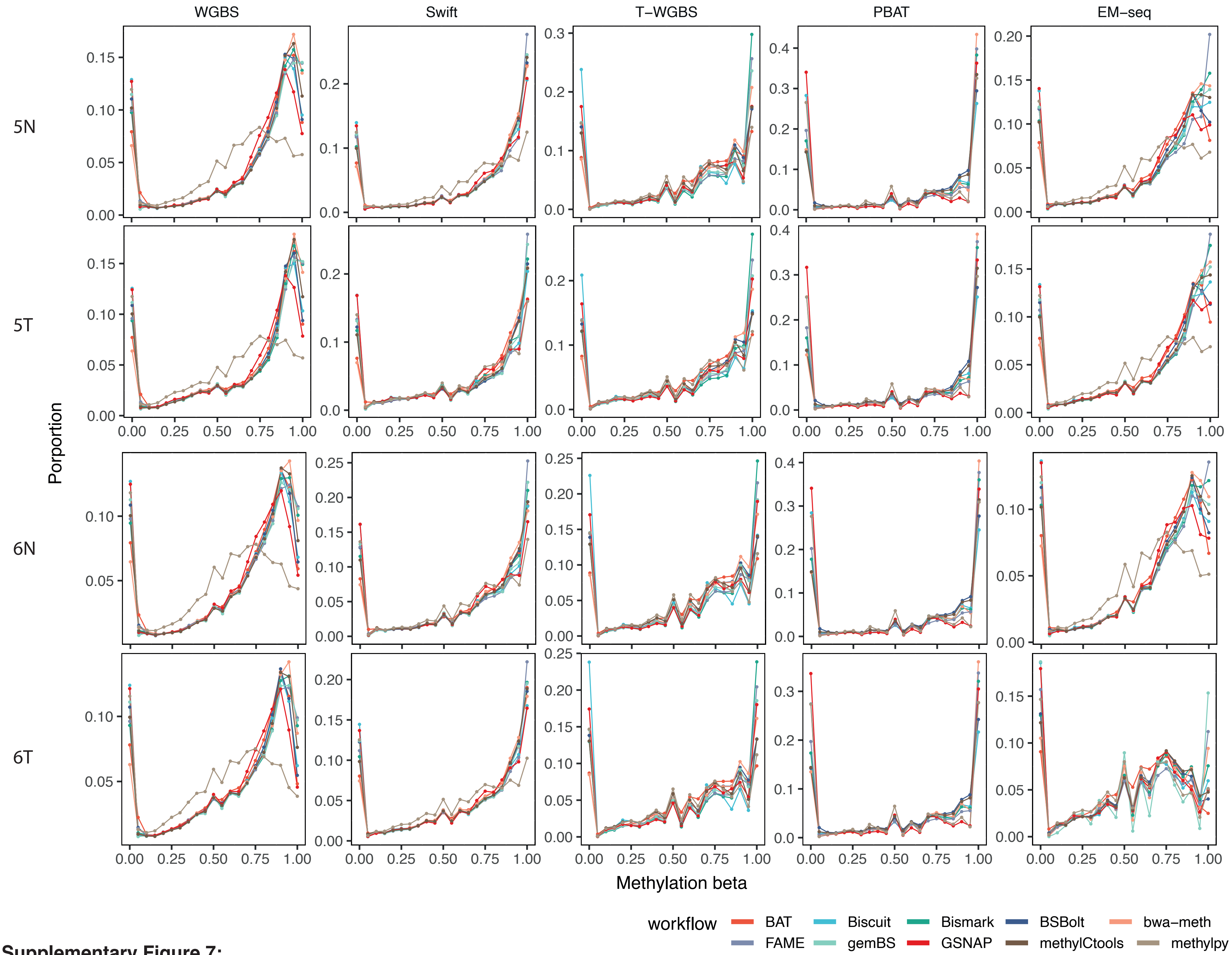


**Supplementary Figure 7:**

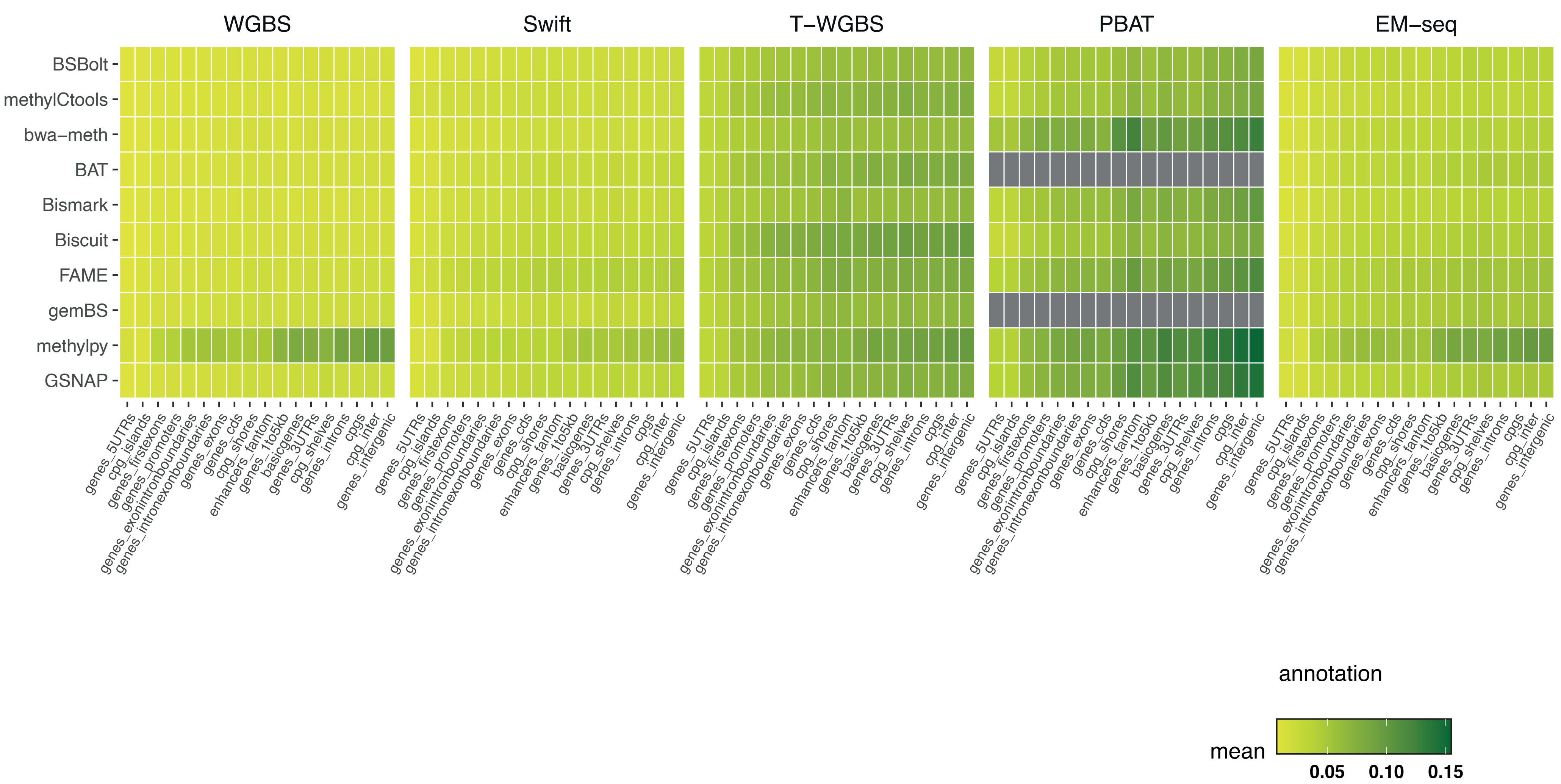


**Supplementary Figure 8:**

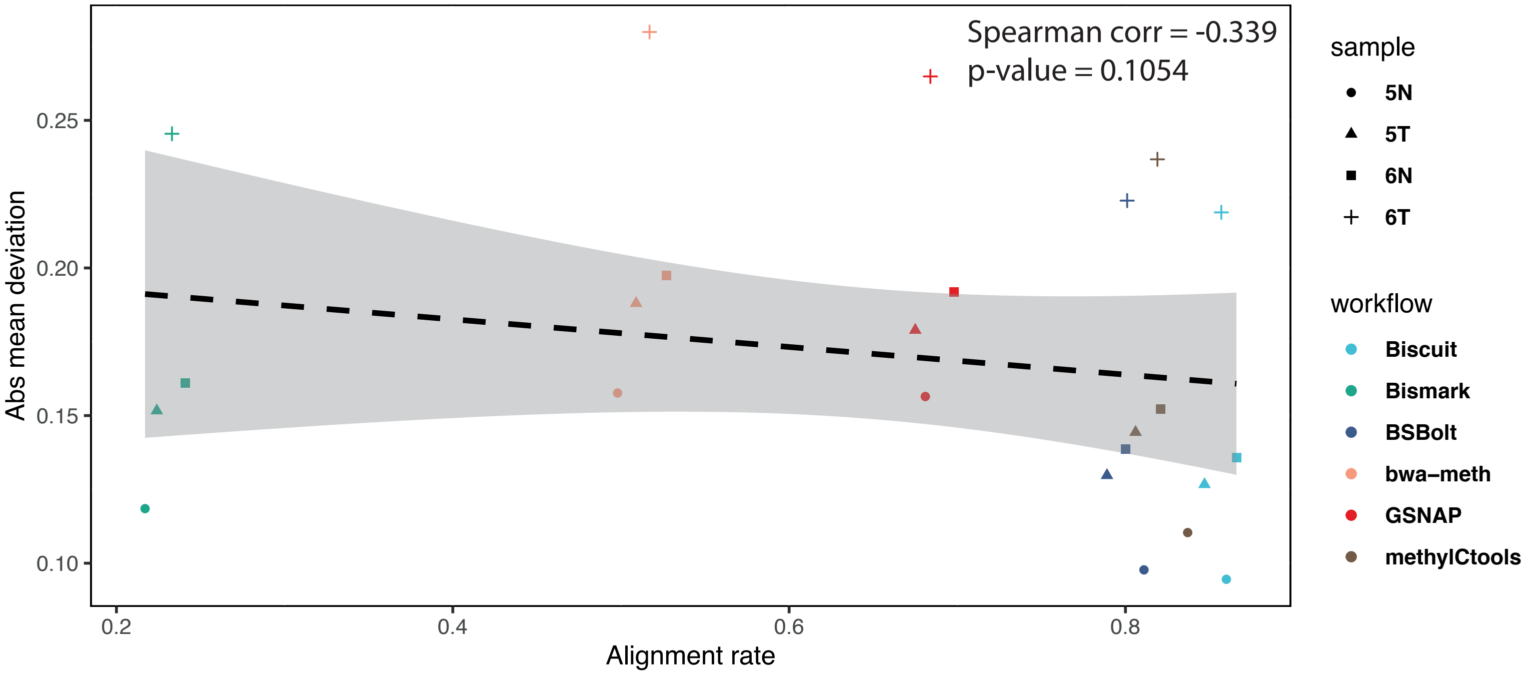


**Supplementary Figure 9:**

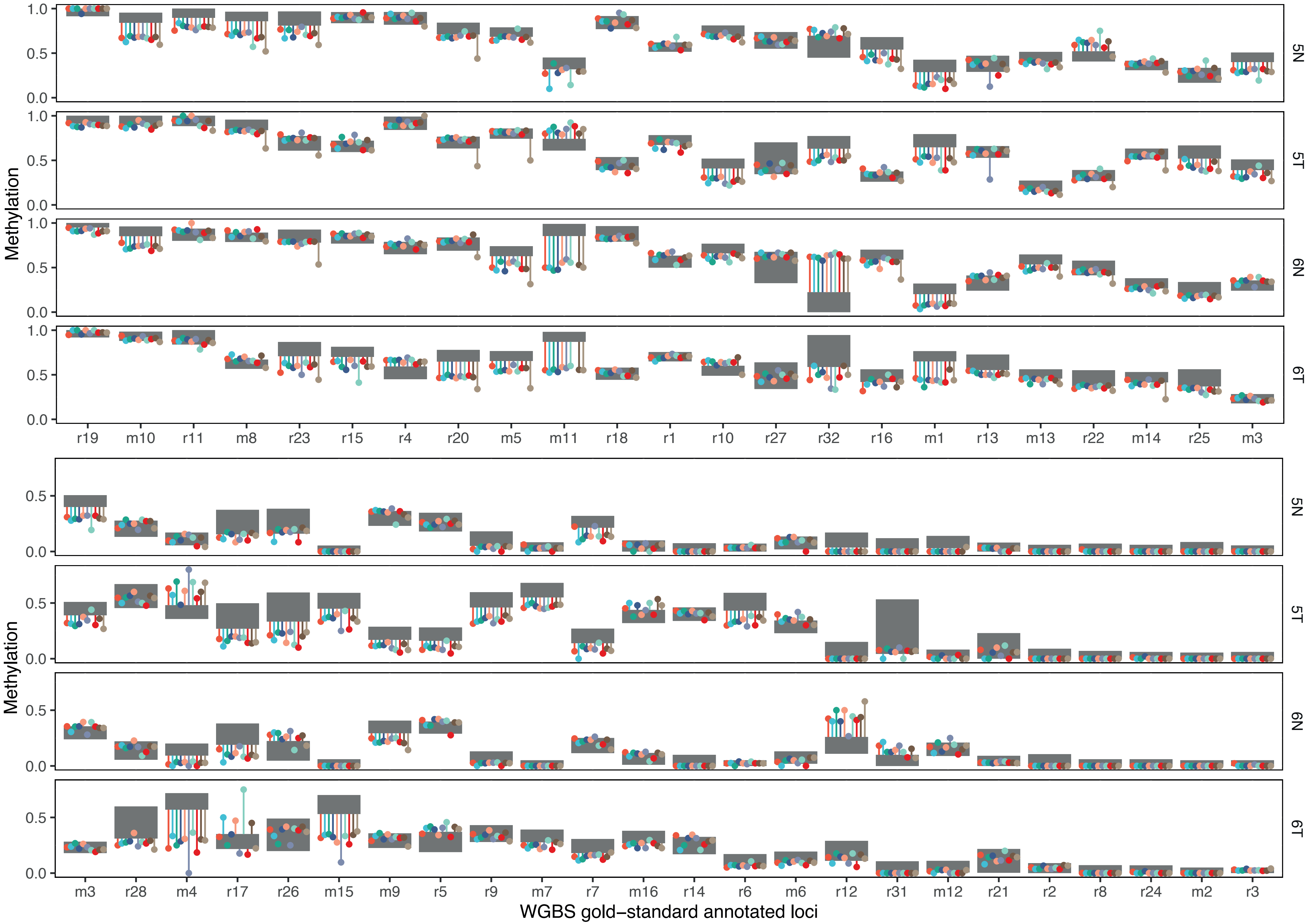


**Supplementary Figure 10:**

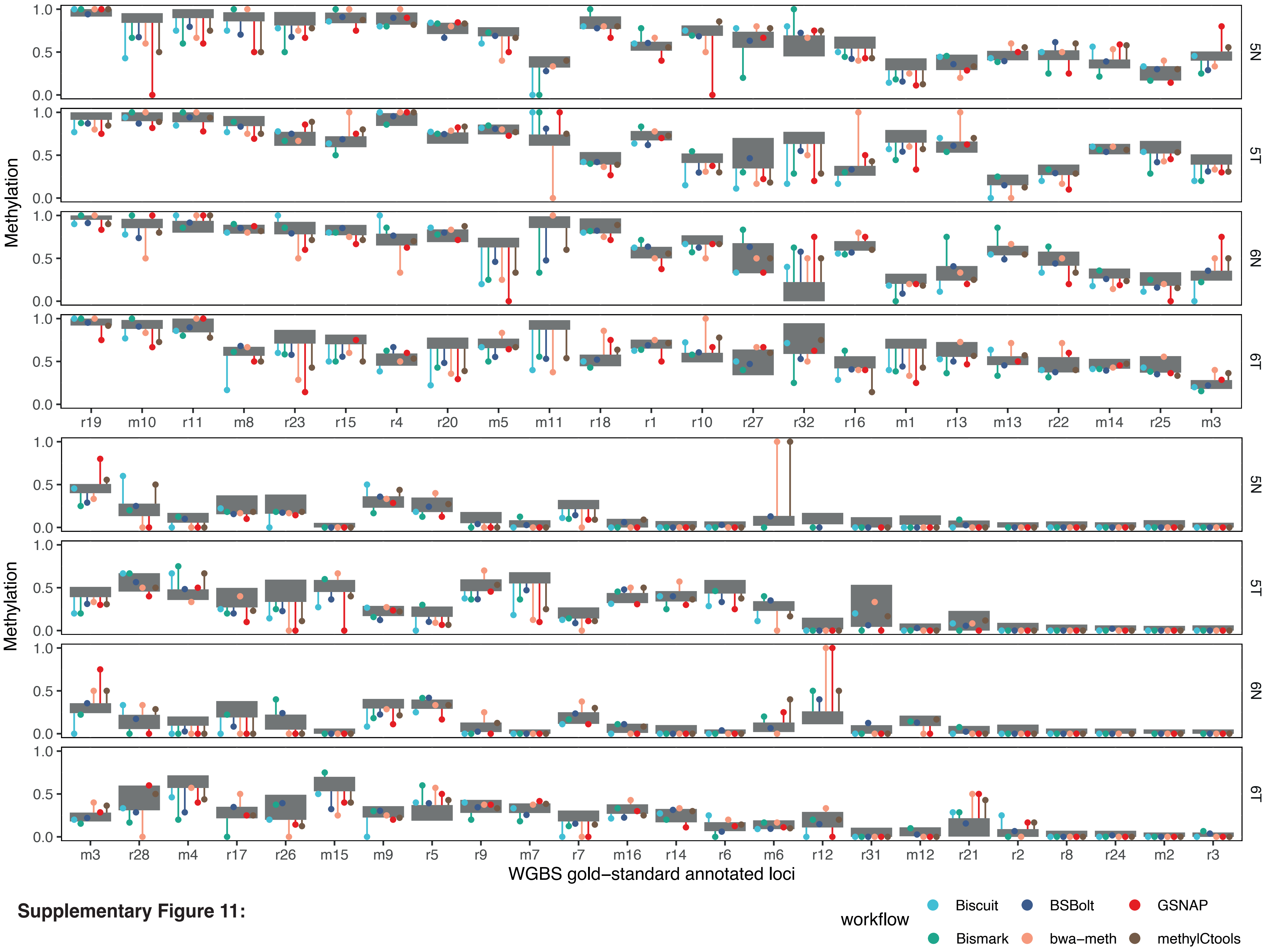


**Supplementary Figure 11:**

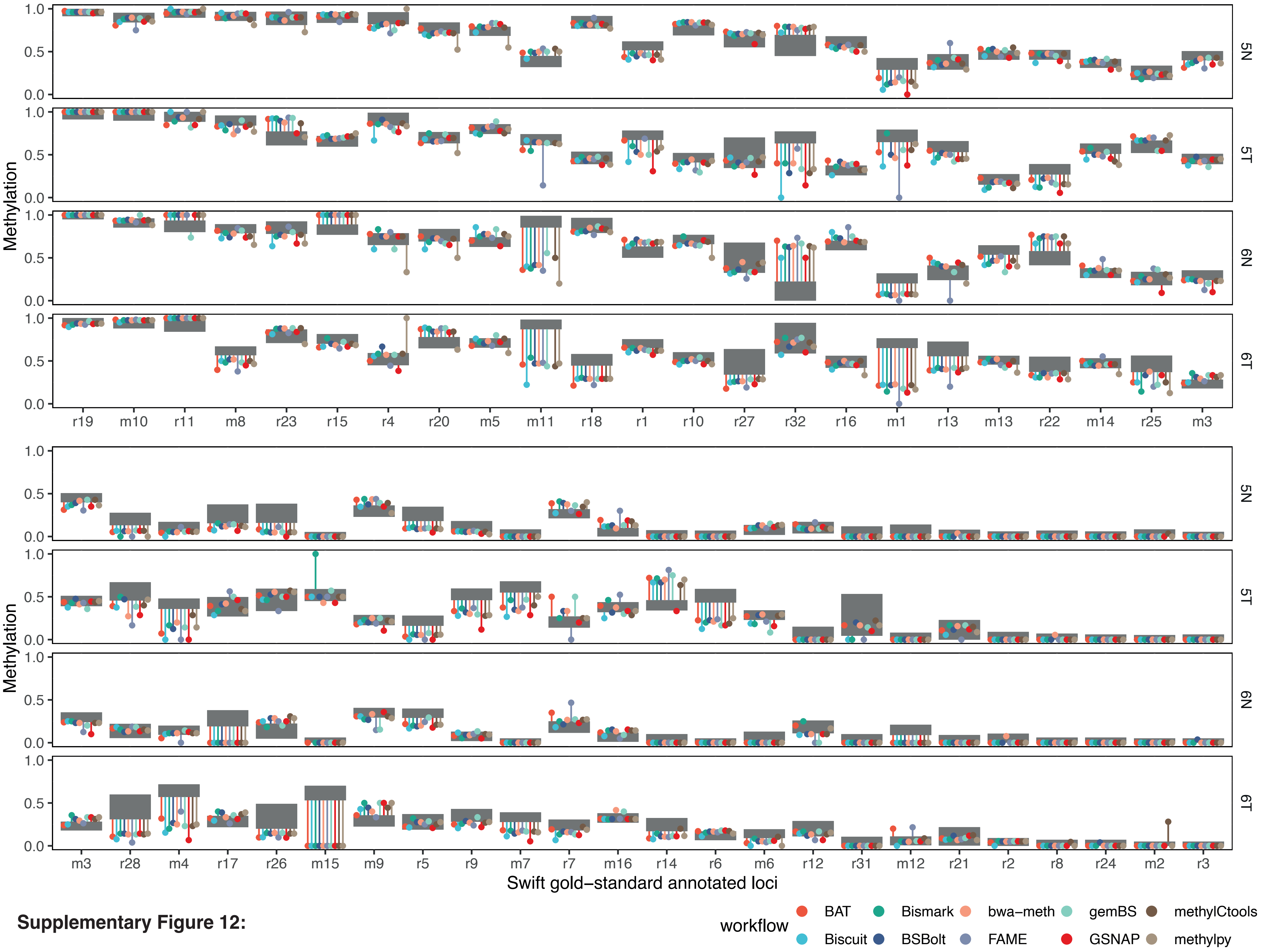


**Supplementary Figure 12:**

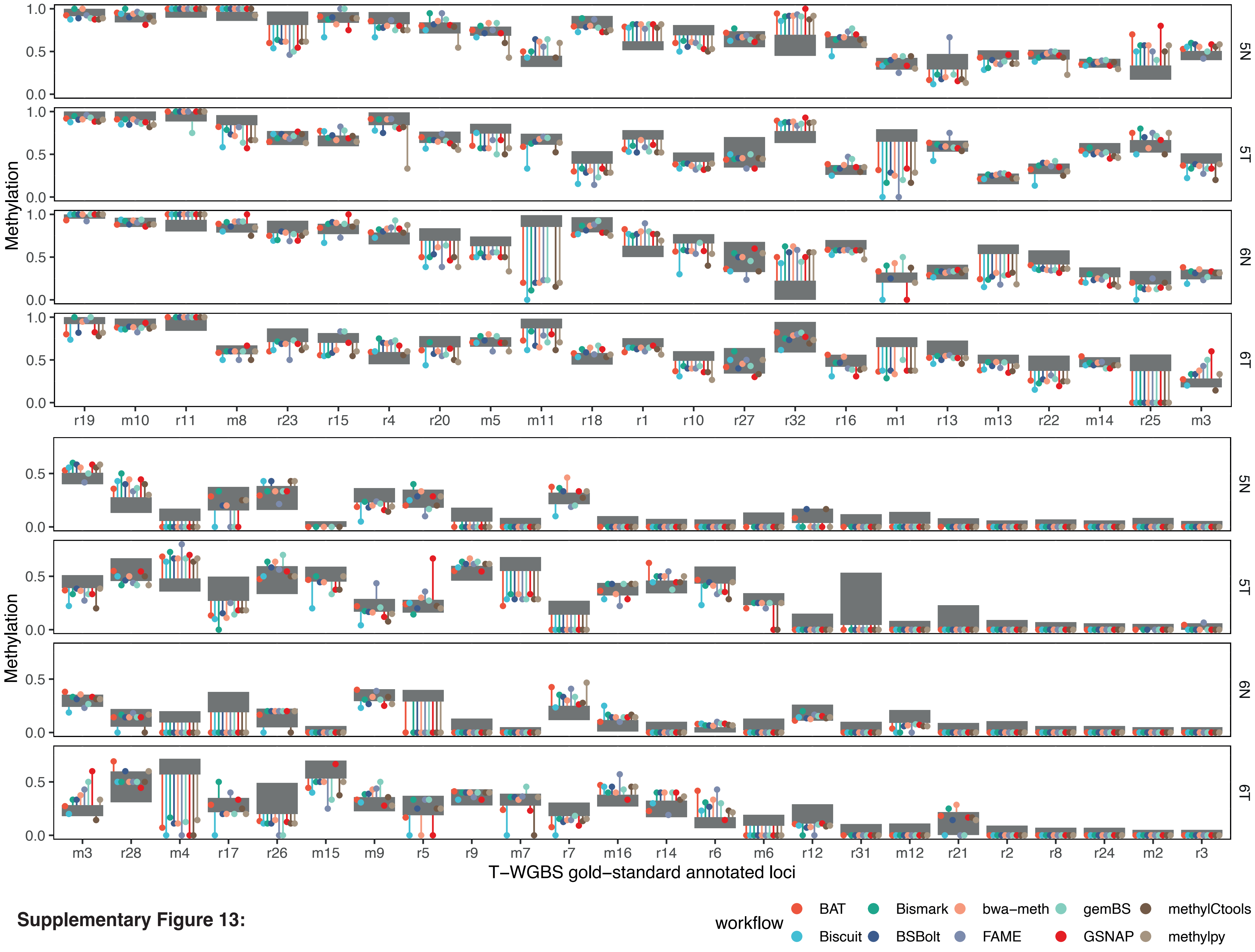


**Supplementary Figure 13:**

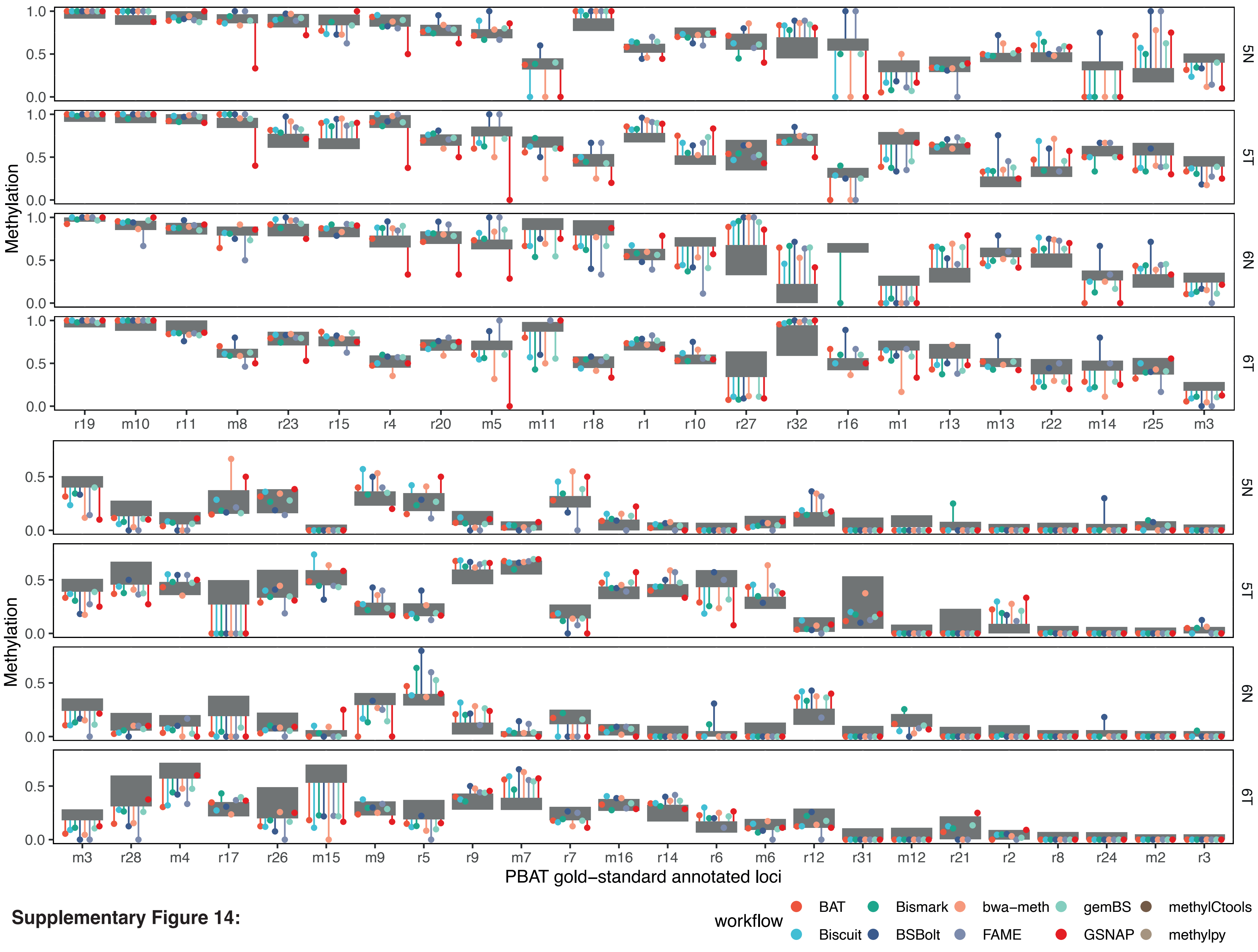


**Supplementary Figure 14:**

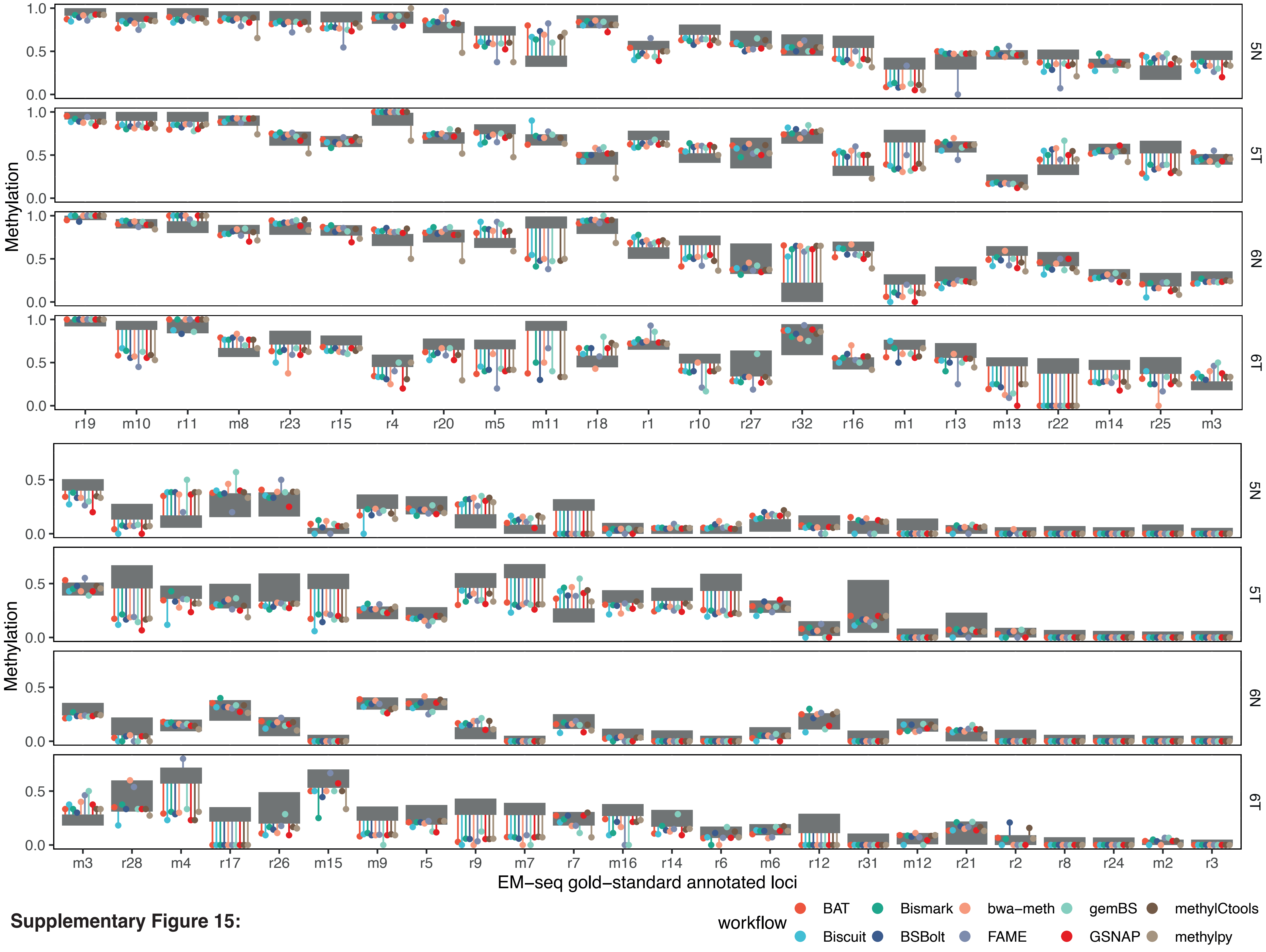


**Supplementary Figure 15:**

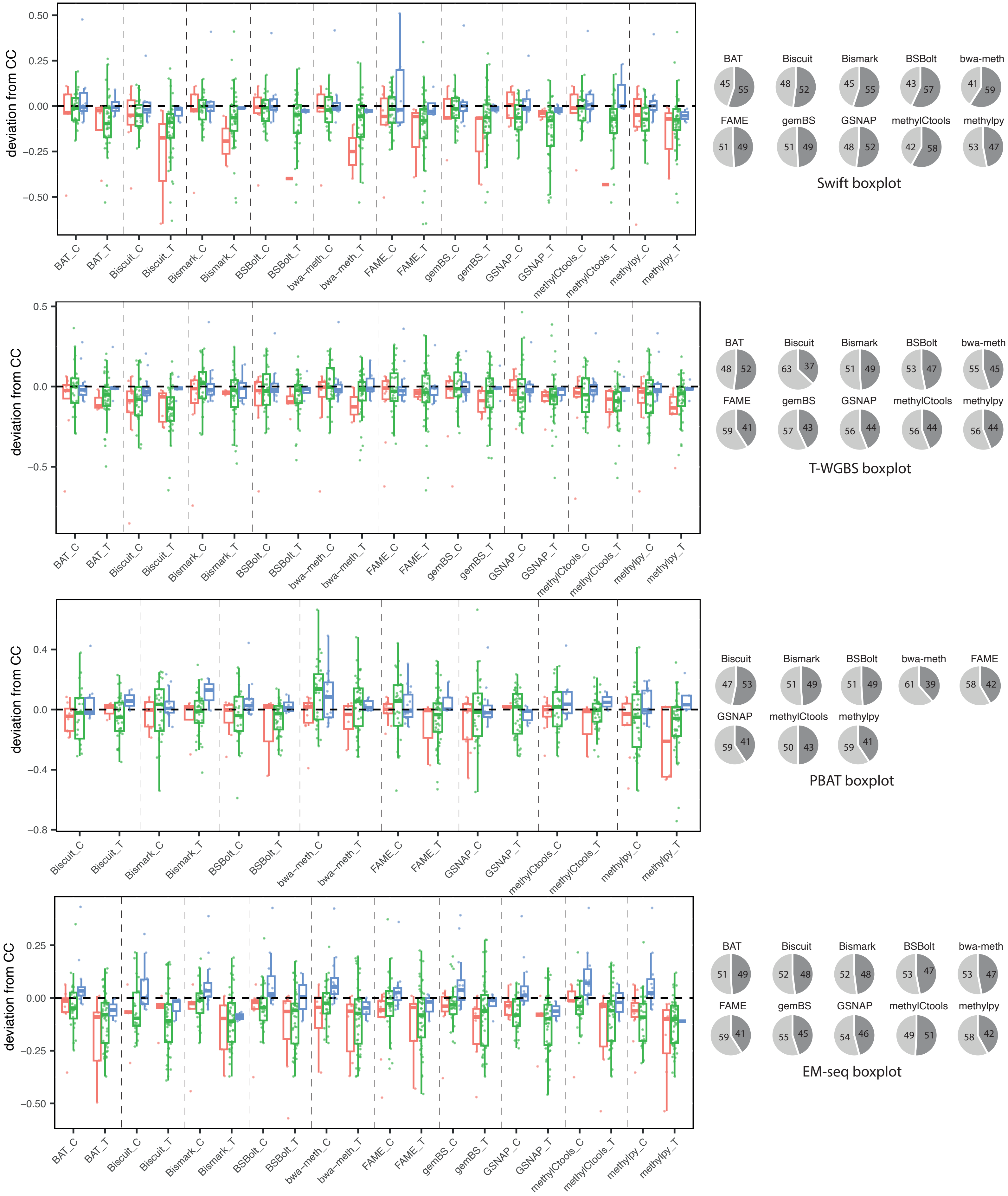


**Supplementary Figure 16:**

a.

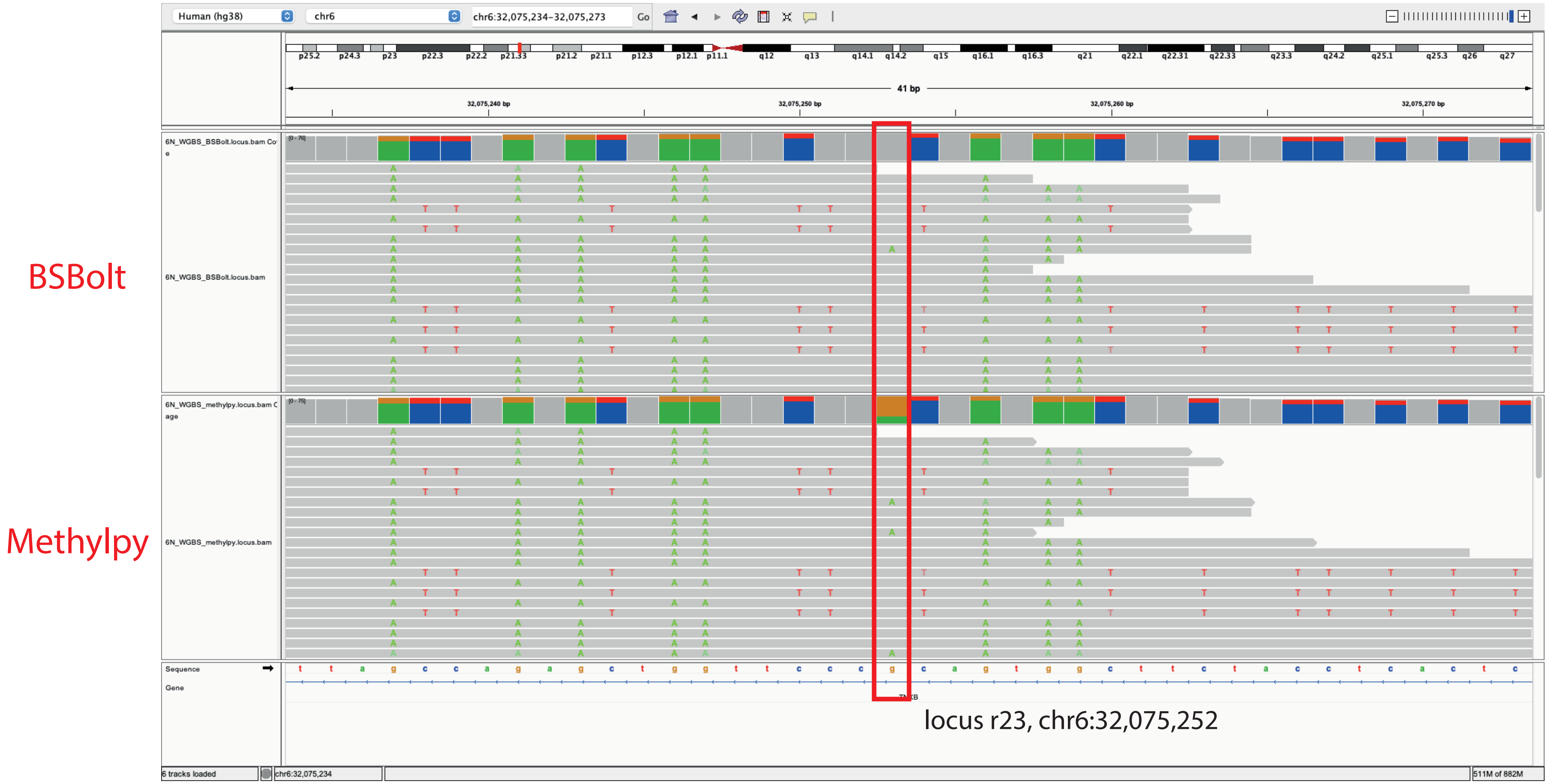


b.

(read name: ST-E00335:86:HWFCLCCXX:6:1211:19229:66409)

```
raw_19229:66409_r1_reverse_complement    ACATTAACCAAAACTAATTCCCGCAATAACTTCTACCTCACTCAAAATAAAATCCAAATC 60
bsbolt_alignment_r1                      ACATTAACCAAAACTAATTCCCGCAATAACTTCTACCTCACTCAAAATAAAATCCAAATC 60
methylpy_intermediate_reverse_r1         ACATTAACCAAAACTAATTCCCACAATAACTTCTACCTCACTCAAAATAAAATCCAAATC 60
methylpy_alignment_r1                    ACATTAACCAAAACTAATTCCCACAATAACTTCTACCTCACTCAAAATAAAATCCAAATC 60
                                         ********************** *************************************
```

**Supplementary Figure 17:**
